# Dual Ribosome Profiling reveals metabolic limitations of cancer and stromal cells in the tumor microenvironment

**DOI:** 10.1101/2025.01.08.631877

**Authors:** Daniela Aviles-Huerta, Del Pizzo Rossella, Alexander Kowar, Ali Hyder Baig, Giuliana Palazzo, Ekaterina Stepanova, Cinthia Claudia Amaya Ramirez, Sara D’Agostino, Edoardo Ratto, Catarina Pechincha, Nora Siefert, Helena Engel, Shangce Du, Silvia Cadenas-De-Miguel, Beiping Miao, Victor Cruz Vilchez, Karin Müller-Decker, Ilaria Elia, Chong Sun, Wilhelm Palm, Fabricio Loayza-Puch

## Abstract

Cancer cells, immune cells, and stromal cells within the tumor microenvironment (TME) collaboratively influence disease progression and therapeutic responses. The nutrient-limited conditions of the TME, particularly the scarcity of glucose, amino acids, and lipids, challenge cancer cell survival^1–4^. However, the metabolic constraints faced by immune and stromal cells in comparison to cancer cells, and how these limitations affect therapeutic outcomes, remain poorly understood. Here, we introduce Dual Ribosome Profiling (DualRP), a method that allows for simultaneous analysis of translation and identification of ribosome stalling, revealing amino acid shortages in different cell types within tumors. Using DualRP, we uncover that interactions between cancer cells and fibroblasts trigger an inflammatory response, mitigating amino acid limitations during glucose starvation. In immunocompetent mouse models, we observe that immune checkpoint blockade therapy induces serine and glycine restrictions specifically in T cells, but not in cancer cells. We further demonstrate that these amino acids are essential for optimal T cell function both *in vitro* and *in vivo*, highlighting their critical role in effective immunotherapy. Our findings show that therapeutic interventions create distinct metabolic demands across different tumor cell types, with nutrient availability significantly influencing the success of immunotherapy. DualRP’s ability to explore cell type-specific metabolic vulnerabilities offers a promising tool for advancing our understanding of tumor biology and improving therapeutic strategies.

## Introduction

Cancers develop in heterogeneous tissue environments, relying on the TME to sustain growth, therapy resistance, and metabolic support^1–3^. In response to nutrient depletion, cancer cells seek additional resources to adapt to a metabolically challenging environment. However, how immune and stromal cell populations respond to nutrient depletion in the TME has been less explored. T cells in particular are very sensitive to this hostile milieu^5–7^. Nutrient scarcity, can severely impede T cell function and response to therapy^8^. While immune checkpoint blockade holds considerable promise, it frequently overlooks the metabolic hurdles that T cells and cancer cells encounter within the TME. Identifying such restrictions in different cells compartments within tumors remains challenging.

Ribosome profiling has emerged as a tool for detecting the availability of amino acids for protein synthesis^9–12^. This approach utilizes global ribosome occupancy to identify limiting amino acids. In principle, critical reduction of intracellular amino acid concentrations leads to tRNA deaminoacylation, resulting in ribosome stalling at a particular codon which is indicative of limitations in the corresponding amino acid. Here, we introduce Dual Ribosome Profiling (DualRP), a method that enables the simultaneous study of translational programs and amino acid restrictions in distinct cell populations, both *in vitro* and *in vivo*. Using DualRP we uncover a metabolic compensatory mechanism triggered by the interaction of cancer cells and fibroblasts and identify specific metabolic vulnerabilities in response to checkpoint inhibition in the T cell compartment of tumors. Our findings provide the rationale to design combinatorial therapies involving check point inhibition and dietary interventions.

## Results

### DualRP enables the study of ribosome occupancy in two interacting cell types

To study metabolic limitations in complex cell populations, we developed DualRP, a technique in which ribosomes from two separate cell populations are labeled with distinct chimeric proteins (GFP-RPL10a, mCherry-RPL10a, or NeonGreen-RPL10a)^13^. This labeling facilitates cell type-specific immunoprecipitation of ribosomes within a mixed population, followed by ribosome profiling (Fig. 1a). To accomplish this in the triple-negative breast cancer (TNBC) cell line SUM-159PT, we introduced N-terminal tags of GFP, mCherry, or mNeonGreen into the *RPL10a* gene through gene editing, achieving homozygous expression (Supplementary Fig 1a-d). Importantly, we verified that chimeric proteins were effectively incorporated into translating ribosomes without adversely affecting cell proliferation or global rates of protein synthesis (Supplementary Fig. 1e-g).

**Figure 1.**
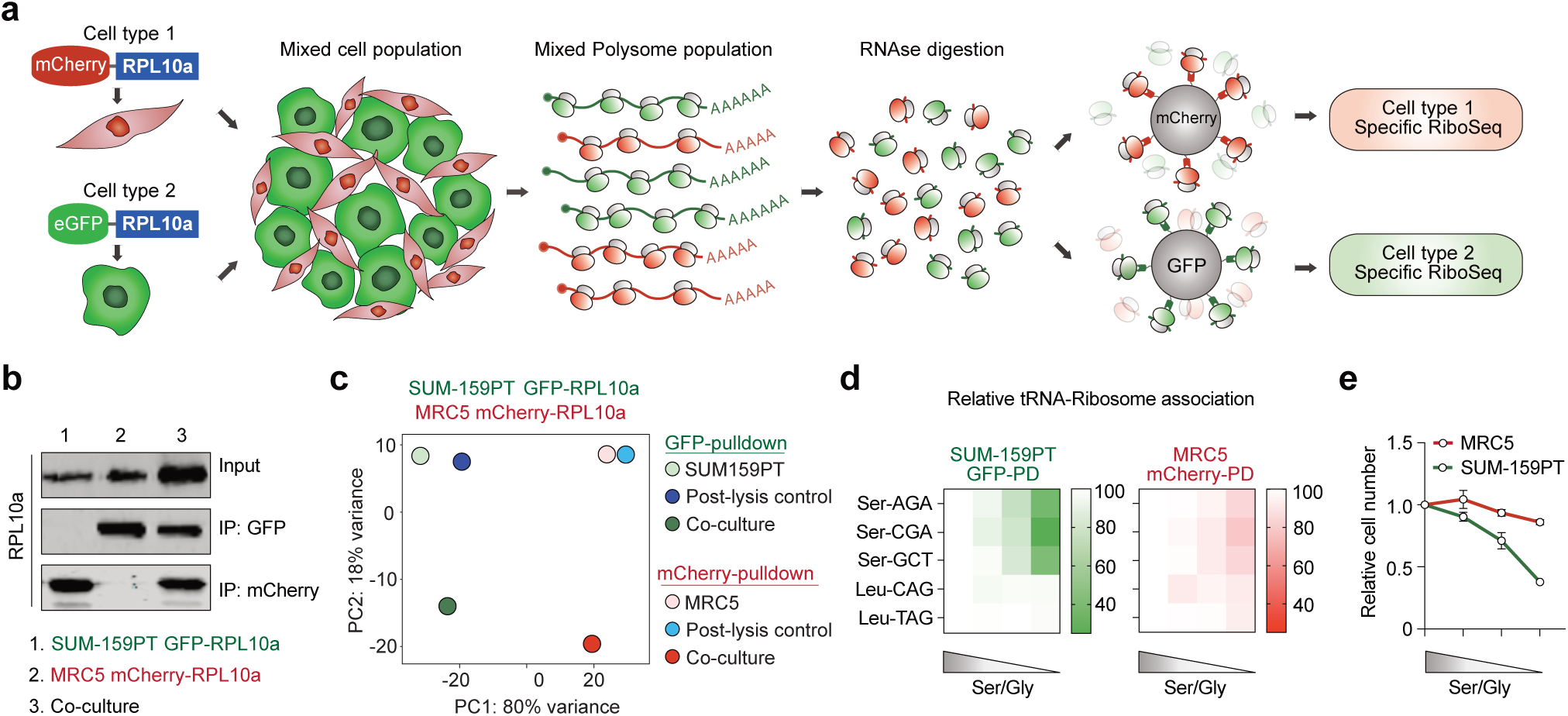
DualRP allows for the study of ribosome occupancy and tRNA-Ribosome association in two interacting cell types. **a,** Schematic diagram of dual ribosome profiling (Dual-RP): Ribosomes from two distinct cell types are individually tagged with chimera proteins, eGFP-RPL10a and mCherry-RPL10a. The two cell populations are mixed, lysed, and ribosomes from each cell type are then immunoprecipitated using highly specific nanobodies. Subsequently, libraries for next-generation sequencing are prepared from the recovered ribosome-protected fragments (RPFs). **b,** Immunoprecipitation experiments with beads coated with nanobodies against GFP (IP: GFP) or against mCherry (IP: mCherry), followed by western blot analysis in SUM-159PT-GFP-RPL10a and MRC5-mCherry-RPL10a cells. **c,** Principal Component Analysis (PCA) from ribosome profiling libraries generated from pull-down of mono- and co-cultures of SUM-159PT-GFP-RPL10a and MRC5-mCherry-RPL10a. Mixes of lysed mono-cultures were used as controls (Post-lysis control). **d,** tRNA-Ribosome association assay in SUM-159PT-GFP-RPL10a and MRC5-mCherry-RPL10a cells starved of serine/glycine (ranging from 0.4mM to 0mM each). PD, pull-down. Data represent mean ± SD (n=3). **e,** Normalized cell numbers at different serine/glycine concentrations. The data are expressed as relative cell numbers compared to the untreated group at the endpoint. Measurements were taken 48 hours after plating. Data represent mean ± SD (n=3).

To assess the effectiveness and quality of the DualRP system, we first generated ribosome profiling libraries from pull-downs of tagged cells and compared them to libraries prepared using conventional sucrose gradients^14^. We observed a strong correlation in ribosome density in all comparisons (Supplementary Fig. 2a-b). Next, we co-cultured SUM-159PT-GFP-RPL10a cells and MRC5-mCherry-RPL10a fibroblasts and performed ribosome immunoprecipitation for each population. Immunoprecipitation assays and principal component analysis (PCA) of RPFs confirmed the specificity of the DualRP approach (Fig. 1b-c). RPFs originating from tagged ribosomes predominantly map to coding sequences (CDSs). These fragments displayed a distinct 3-nt periodicity and exhibited a heightened density at the start codon (Supplementary Fig. 2c-f). Our results indicate that DualRP enables highly consistent and high-quality ribosome occupancy measurements in two interacting populations.

Amino acid starvation results in deaminoacylation of the corresponding tRNAs, leading to reduced interactions with site A of the ribosome^15,16^. DualRP can also be used to assess the interactions between tRNAs and ribosomes (Supplementary Fig. 2g). Through the pull-down of tagged ribosomes in co-cultures of breast cancer cells and fibroblasts, we observed that cancer cell ribosomes exhibited decreased association with tRNAs when subjected to limiting concentrations of glutamine or serine/glycine (Fig. 1d; Supplementary Fig. 2h-j). Importantly, aminoacylation rates in both cell populations were strongly correlated with cell growth (Fig. 1e; Supplementary Fig. 2h-j). In summary, DualRP offers the unique ability to simultaneously investigate ribosome occupancy and the association between ribosomes and tRNAs in two interacting populations.

### DualRP elucidates a type I IFN-mediated pathway that enhances amino acid availability for tRNA charging

To understand how different types of interacting cells respond to metabolically challenging conditions, we exposed co-cultures of breast cancer cells (SUM-159PT-GFP-RPL10a) and fibroblasts (MRC5-mCherry-RPL10a) to limiting concentrations of glucose. Using DualRP and differential ribosome codon reading (diricore) analysis^9^, we discovered that cancer cells experienced ribosome stalling at alanine, glycine, and valine codons when grown individually as monocultures, indicating limitations in the corresponding amino acids (Fig. 2a). However, alanine and glycine deficiencies were remarkably alleviated when the cells were co-cultured. (Fig. 2a). The same effect was observed in MDA-MB-231-GFP-RPL10a cells co-cultured with MRC5 fibroblasts (mCherry-RPL10a) and MDA-MB-231 cells carrying a different tag (NeonGreen-RPL10a; Supplementary Figs. 3a and 3d). Additionally, a similar response was noted in SUM-159PT cells where the RPS3 gene was C-terminally tagged with GFP (RPS3-GFP; Supplementary Fig. 1h-l), indicating that ribosome stalling during glucose starvation is independent of the cell line, protein tag, or ribosomal subunit tagging (Supplementary Fig. 3g). These findings suggest that the interplay between cancer cells and fibroblasts facilitate the exchange or synthesis of nutrients, providing a mechanism by which these cells collaboratively overcome the amino acid constraints imposed by glucose limitation.

**Figure 2.**
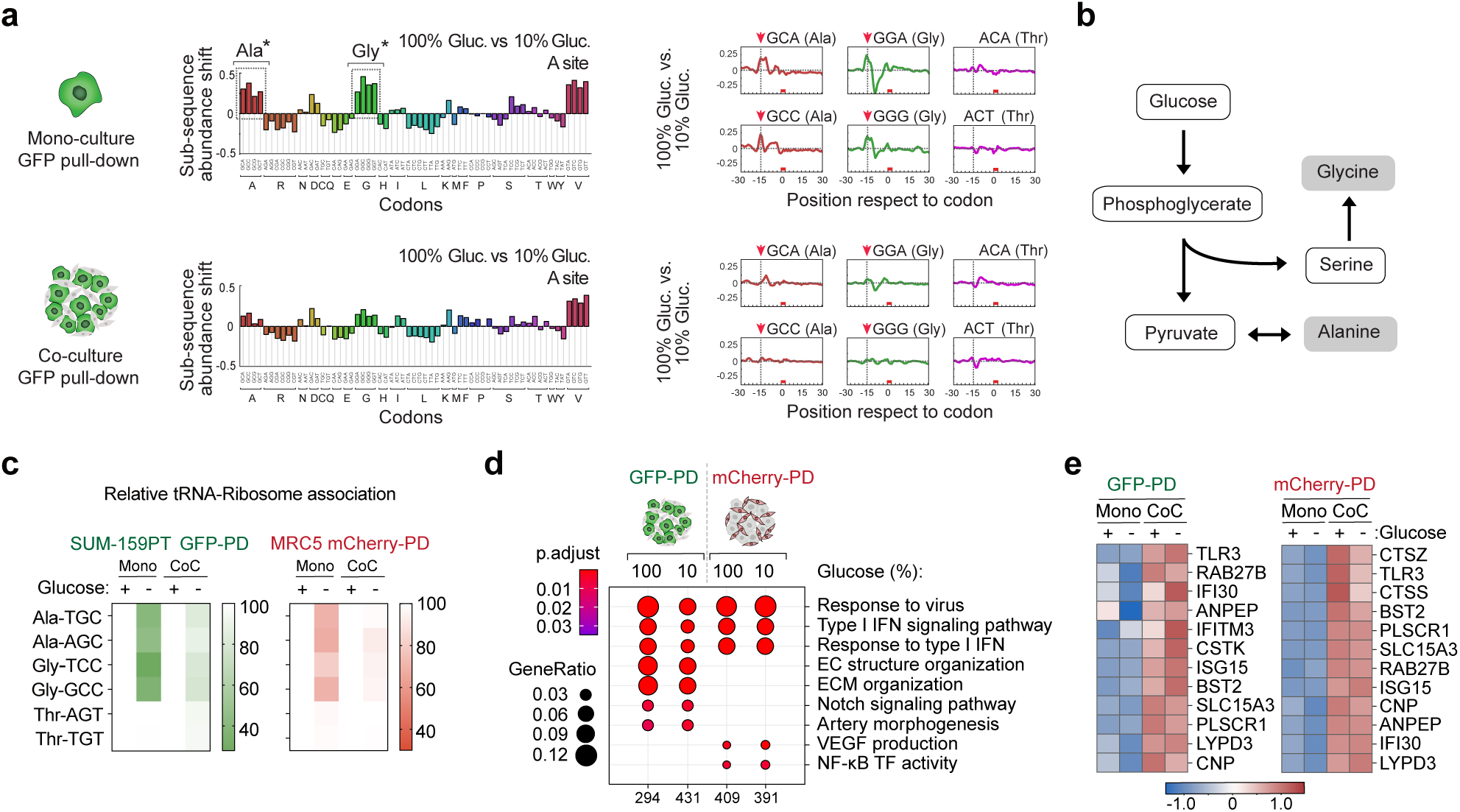
DualRP reveals that heterotypic cell interactions restore ribosome stalling and tRNA aminoacylation rates during glucose starvation. **a,** Diricore analysis of GFP pull-downs from mono-cultures of SUM-159PT GFP-RPL10a cells (upper panel) and co-cultures with MRC5 mCherry-RPL10a (lower panel) grown in full (100%, 25mM) or glucose-deprived (10%, 2,5 mM) medium. Cells were treated for 48 hrs. The dip observed at position -12 in the ribosome density plots (right panels) reflects a reduced ribosome density at the P site of the codon. *Out-of-frame analysis P < 0.01 for the GCA, GCC, GCG, GCT Ala codons and for the GGA, GGC, GGG Gly codons. **b,** Schematic diagram of the synthesis of the amino acids alanine and glycine from glucose. **c,** tRNA-Ribosome association assay in mono- and co-cultures (CoC) of SUM-159PT-GFP-RPL10a and MRC5-mCherry-RPL10a growing in full ((+), 25mM) or glucose-deprived ((-), 2,5mM) medium. Data represent mean ± SD (n=3). PD, pull-down. **d,** Gene ontology analysis of genes upregulated upon heterotypic interaction of SUM-159PT GFP-RPL10a and MRC5 mCherry-RPL10a growing in full (100%, 25mM) or glucose-deprived (10%, 2,5mM) medium. Significantly enriched gene sets (p-value < 0.05) are shown, along with the number of identified proteins in the respective gene set (GeneRatio). PD, pull-down. **e,** Heatmaps displaying differential expression of representative lysosomal genes based on Ribo-seq counts in mono- and co-cultures of SUM-159PT GFP-RPL10a and MRC5 mCherry-RPL10a grown in full ((+), 25mM) or glucose-deprived ((-), 2,5mM) medium. LogFC is shown; PD, pull-down.

Alanine and glycine can be generated *de novo* from glucose through pyruvate transamination and the serine synthesis pathway, respectively^17,18^ (Fig. 2b). TNBC cells lacking PHGDH rely heavily on extracellular serine, which can be taken up from the environment or synthesized from glycine by serine hydroxymethyltransferase (SHMT)^19^. This glycine-to-serine conversion becomes crucial when glucose-derived *de novo* synthesis is impaired. Consistent with ribosome stalling, we observed a decrease in ribosome association with aminoacylated alanine (Ala) tRNAs TGC and AGC, as well as glycine (Gly) tRNAs TCC and GCC, after glucose starvation in monocultures of cancer cells and fibroblasts. By contrast, the association of aminoacylated control threonine (Thr) tRNAs AGT and TGT remained unaffected (Fig. 2c). However, in glucose-deprived co-cultures of SUM-159PT cells and MRC5 fibroblasts, the rate of ribosome association with Ala- and Gly-tRNAs was restored (Fig. 2c). A similar pattern was observed in MDA-MB-231, SUM-159PT-NeonGreen-RPL10a, and SUM-159PT-RPS3-GFP cells when co-cultured with MRC5 fibroblasts (Supplementary Figs. 3b, 3e, and 3h). These results collectively confirm the effectiveness of the DualRP system in investigating cell-type-specific amino acid deficiencies within cultured cells and reveal that the heterotypic cell interactions restore tRNA aminoacylation rates during glucose starvation.

To explore the mechanism behind the restoration of tRNA aminoacylation in metabolically challenged co-cultures, we examined differences in Ribo-Seq-based expression signatures between mono- and co-cultures of breast cancer cells and fibroblasts in both full media and under limiting concentrations of glucose. Gene ontology (GO) and gene set enrichment analysis (GSEA) highlighted the significant enrichment of terms related to the type I IFN signaling pathway in both cell lines (Fig. 2d). Specific terms in SUM-159PT cells were associated with extracellular matrix (ECM) organization and the Notch signaling pathway, while MRC5 cells in co-culture upregulated genes involved in VEGF production and NF-κB transcription factor activity (Fig. 2d). These expression patterns were also observed in co-cultures grown with limiting concentrations of glucose in different cell lines and with different ribosomal tags (Supplementary Fig. 3c, 3f, and 3i).

Among metabolic gene subsets, we observed a robust increase in the expression of genes localized in the lysosomes of both breast cancer cells and MRC5 fibroblasts (Fig. 2e; Supplementary Fig. 4a-e). The promoters of the upregulated lysosomal genes were found to be enriched in STAT1 binding sites, indicating their potential as interferon-stimulated genes (ISGs) (Supplementary Fig. 4f). As these co-cultures induced the IFN-I signaling pathway, activating JAK-STAT signaling and the expression of ISGs^20^, this observation suggested that the subset of lysosomal genes might directly respond to IFN-I signaling pathway activation, thereby enhancing lysosomal nutrient generation.

To investigate this possibility, we measured lysosomal proteolysis using DQ BSA, a self-quenched albumin probe whose fluorescence becomes dequenched upon lysosomal proteolysis^21^ (Fig. 3a). Co-culturing resulted in a significant increase in intracellular DQ BSA fluorescence in both breast cancer cells and fibroblasts (Fig. 3b), indicating enhanced lysosomal catabolism. To determine whether this response was mediated by the IFN-I signaling pathway or glucose starvation, we treated mono-cultures of SUM-159PT cells with either IFN-β or limiting concentrations of glucose. Glucose depletion alone did not enhance lysosomal catabolism, whereas IFN-β treatment increased DQ BSA fluorescence both in full medium and under glucose-limiting conditions (Supplementary Fig. 4g). Next, we generated STAT1 knockout SUM-159PT cells (Supplementary Fig. 4h). Upon co-culture, these knockout cells failed to upregulate canonical ISGs and genes associated with lysosomal function (Supplementary Fig. 4i). Furthermore, their intracellular DQ BSA fluorescence did not increase, either in full medium (Figs. 3c-d) or under glucose starvation (Supplementary Fig. 4j). STAT1 knockout cells were also unable to rescue the ribosome association of Ala- and Gly-tRNAs in glucose-deprived co-cultures (Fig. 3e). Thus, the IFN-I signaling pathway is required to promote lysosomal function and amino acid production in co-cultures exposed to limiting nutrient concentrations.

**Figure 3.**
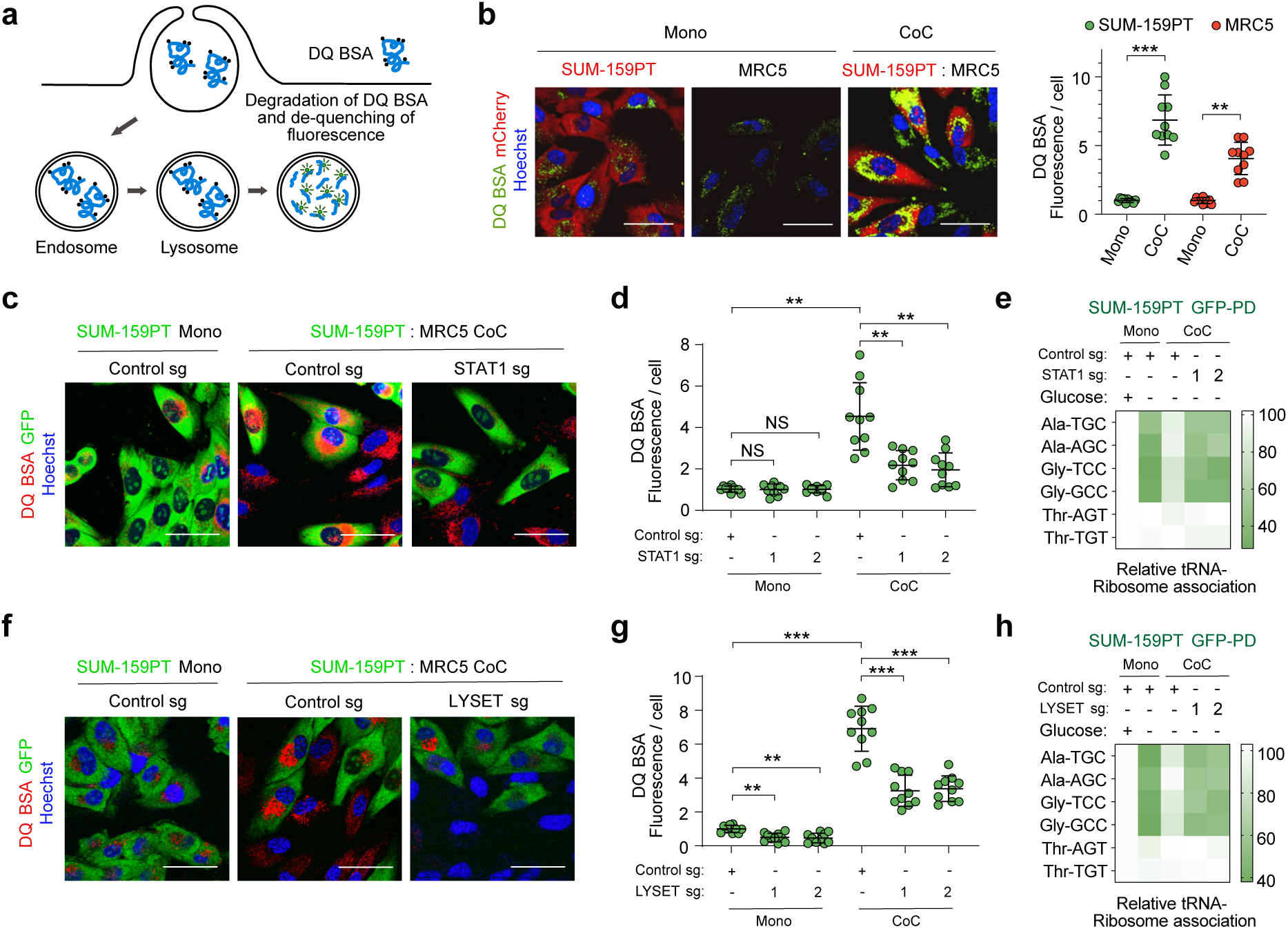
Lysosomal catabolism is enhanced upon heterotypic interactions in a type I IFN-dependent manner. **a,** Schematic showing endo-lysosome formation and DQ BSA degradation. DQ BSA undergoes cellular endocytosis and transits from early endosomes to late endosomes. These late endosomes subsequently merge with lysosomes rich in acidic proteases. As a result, endo-lysosomes form, facilitating the degradation of DQ BSA and consequently restoring the fluorescence of the dye associated with this cargo. **b,** Representative fluorescence microscopy images showing the degradation of lysosomal DQ BSA in mono- and co-cultures (CoC) of SUM-159PT-mCherry-RPL10a and MRC5 cells. Scale bars = 50 µm. Quantification of DQ BSA fluorescence in the indicated conditions (Right panel). Data represent mean ± SD (n ≥10); **P < 0.01; ***P < 0.001 by Student’s t-test. **c,** Representative confocal microscopy images illustrating the degradation of lysosomal DQ BSA in SUM-159PT-GFP-RPL10a expressing sgRNAs targeting the STAT1 and growing as mono- or co-cultures with MRC5. Scale bars = 50 µm. **d,** Quantification of DQ BSA fluorescence in SUM-159PT GFP-RPL10a transduced with sgRNAs targeting the STAT1 gene or a control sequence growing as mono- or co-cultures (CoC) with MRC5 cells. The cells were maintained in full medium (Glucose 25mM). Data represent mean ± SD (n ≥10); **P < 0.01; ***P < 0.001 by Student’s t-test. **e,** Quantification of tRNA-Ribosome association in SUM-159PT-GFP-RPL10a expressing sgRNAs targeting STAT1 grown as mono- or co-cultures (CoC) with MRC5 cells in full ((+), 25mM) or glucose-deprived ((-), 2,5mM) medium. Data represent mean ± SD (n=3) **f,** Representative confocal microscopy images showing the degradation of lysosomal DQ BSA in SUM-159PT GFP-RPL10a expressing sgRNAs against LYSET and growing as mono- or co-cultures with MRC5 cells. Scale bars = 50 µm. **g,** DQ BSA quantification in SUM-159PT GFP-RPL10a transduced with sgRNAs targeting the LYSET gene growing as mono- or co-cultures with MRC5 cells. Cells were grown in full medium (Glucose 25mM). Data represent mean ± SD (n ≥10); **P < 0.01; ***P < 0.001 by Student’s t-test. **h,** tRNA-Ribosome association assay in SUM-159PT-GFP-RPL10a expressing sgRNAs against LYSET growing as mono- or co-cultures with MRC5 cells in in full ((+), 25mM) or glucose-deprived ((-), 2,5mM) medium. Data represent mean ± SD (n=3). PD, pull-down.

Lysosomal degradation of extracellular proteins serves as an amino acid source that cancer cells exploit to thrive under nutrient-poor conditions^22,23^. To establish whether lysosomal catabolism is necessary for producing charged tRNAs in glucose-starved co-cultures, we generated breast cancer cells deficient for LYSET (Supplementary Fig. 4k), which was recently identified to be required for lysosomal catabolism of macropinocytic and autophagic cargoes^24^. LYSET KO cells displayed reduced enzymatic activities of various lysosomal proteases but exhibited induction of ISGs to the same extent as control cells upon IFN-β treatment (Supplementary Fig. 4l-m). DQ BSA fluorescence and ribosome association of Ala- and Gly-tRNAs were strongly decreased in LYSET-deficient cells upon co-culture with MRC5 cells (Fig. 3f-h; Supplementary Fig. 4k). Thus, the interaction between breast cancer cells and stromal cells enhances lysosomal catabolism, ultimately leading to greater nutrient availability for tRNA aminoacylation.

### DualRP detects specific metabolic limitations in different cell compartments of the TME

Having established the DualRP system in cells in culture, we sought to identify metabolic constraints simultaneously in distinct cell types within tumors. For this purpose, we adapted DualRP for studying amino acid restrictions in the cancer and T cell compartments of the TME. We crossed mice expressing the hemagglutinin (HA)-tagged ribosomal protein RPL22 (RiboTag mice) with CD4-Cre mice to selectively tag ribosomes in T cells with HA (CD4-Cre:RiboTag mice; Supplementary Fig. 5a)^25^. Simultaneously, we engineered a mouse tumor cell line (E0771) derived from spontaneous breast cancer in C57BL/6 mice, where the endogenous *Rpl10a* gene was homozygously tagged at the N-terminus with GFP (Supplementary Fig. 5b-d). We confirmed that GFP-RPL10a-expressing cells did not show proliferative or translational defects (Supplementary Fig. 5e-g). Western blot analysis of proteins immunoprecipitated with anti-HA from CD4-Cre:RiboTag homozygous mouse spleens demonstrated that RPL22-HA was exclusively detectable in mouse T cells and co-immunoprecipitated with other ribosomal proteins (RPL10a). By contrast, anti-HA did not pull-down ribosomal proteins from E0771-GFP-RPL10a cells (Supplementary Fig. 5h), confirming the specificity of our approach.

To enable *in vivo* interactions between tumor cells and host-derived T cells, we orthotopically injected CD4-Cre:RiboTag mice with E0771-GFP-RPL10a cells (Supplementary Fig. 6a). After tumors reached a defined size and recruited T cells, we collected tumor samples, flash-frozen them to capture native gene expression patterns and ribosome positions, and subsequently conducted DualRP. RPFs derived from both cancer and T cells predominantly mapped to coding sequences (CDSs), exhibiting a clear 3-nt periodicity and displaying an increased density at the start codon (Supplementary Fig 5.i-m). Principal component analysis (PCA) of RPFs confirmed the specificity of DualRP in tumors (Supplementary Fig. 5n). Thus, DualRP can be used to study ribosome occupancy in multiple tumor cell compartments with high specificity.

To identify metabolic constraints within the cancer and T cell compartments in response to immune checkpoint blockade therapy, we administered anti-PD1 (αPD1) to CD4-Cre:RiboTag tumor-bearing mice and conducted DualRP analysis (Fig. 4a). Ribosome occupancy analysis revealed ribosome stalling at alanine, aspartic acid, serine, and glycine codons within the T cell compartment, while cancer cells showed no enrichment in any specific codons (Fig. 4b; Supplementary Fig. 6b). These findings suggest that these amino acids may be essential for an effective T cell cytotoxic response.

**Figure 4.**
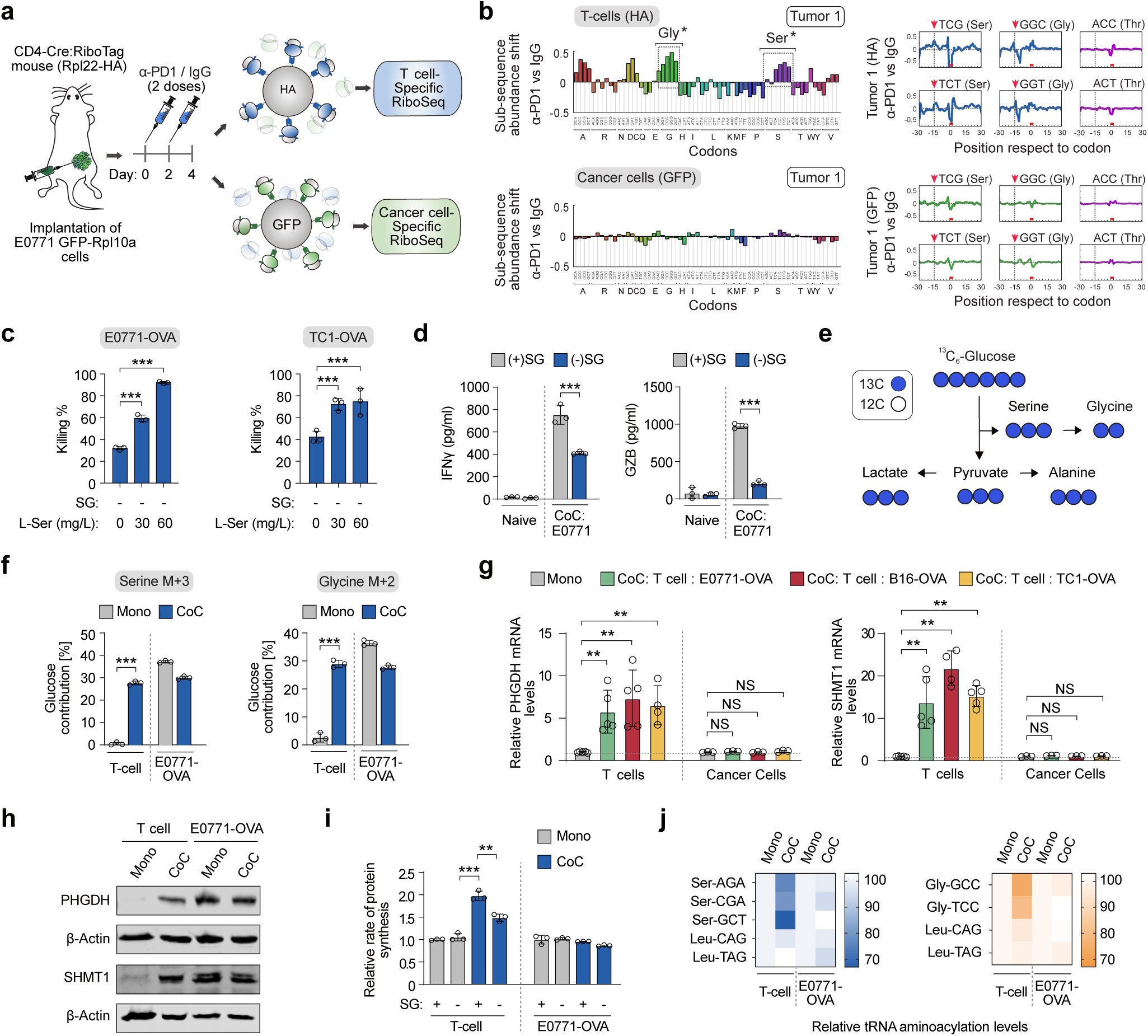
Serine and glycine are limiting amino acids in the T cell compartment of tumors and are necessary for an effective T cell cytotoxic response. **a**, Schematic of experimental design. Mice were treated with two doses of anti-PD1(2 µg/µl) or IgG (2 µg/µl). Two days after the last dose, tumors were flash frozen, lysed, and ribosomes from each cell type were immunoprecipitated using either anti-HA or anti-GFPs coated beads. **b,** Diricore analysis of tumors treated as described in (a). Ribosome stalling in tumor-infiltrated T cells and E0771 tumor cells are shown in the upper and lower panels, respectively. Ribosome density at the A site is shown. *Out-of-frame analysis P < 0.01 for the TCA, TCC, TCG, TCT Ser codons and for the GGA, GGC, GGG Gly codons. The dip observed at position -12 in the ribosome density plots (right panels) reflects a reduced ribosome density at the P site of the codon. **c,** Killing assay by CD8+ T cells in co-culture with E0771-OVA or TC1-OVA cells in serine/glycine (SG)-deprived medium. Data represent mean ± SD (n = 3); ***P < 0.001 by Student’s t-test. **d,** Quantification of IFNγ and GzmB levels produced by CD8+ T cells when co-culture with E0771-OVA cancer cells in full medium or SG-deprived medium. Data represent mean ± SD (n = 3); ***P < 0.001 by Student’s t-test. **e,** Schematic for ^13^C_6_-Glucose tracing into the serine/glycine synthesis pathway. Blue circles represent ^13^C atoms while white circles represent ^12^C atoms. **f,** Contribution of ^13^C_6_-Glucose to serine M+3 and glycine M+2 in OT-I CD8+ T cells and E0771-OVA breast cancer cells growing as mono- or co-cultures. Data represent mean ± SD (n = 3); ***P < 0.001 by Student’s t-test. **g-h,** qRT-PCR quantification (g) and Western blotting (h) of the indicated genes in co-cultures of OT-I CD8+ T cells and OVA-expressing cancer cell lines. Cells were separated by size after 24 hours of co-culture (Supplementary Fig. 6c). Data represent mean ± SD (n = 5 for T cells and n=3 for cancer cells); NS, not significant; **P < 0.01; by Student’s t-test. **i,** Protein synthesis rates based on OP-Puro incorporation in OT-I CD8+ T cells and E0771-OVA breast cancer cells growing as mono- or co-cultures. Cells were grown in full medium (SG+) or serine/glycine-deprived medium (SG-). Data represent mean ± SD (n = 3); **P < 0.01; ***P < 0.001 by Student’s t-test. **j,** Ser-tRNAs, Gly-tRNAs and control Leu-tRNAs aminoacylation analysis in OT-I CD8+ T cells and E0771-OVA breast cancer cells growing as mono- or co-cultures.

Given that serine and glycine can be interconverted—and serine is particularly critical for T cell activation^26^—we focused on these two amino acids. To further validate these observations, we used a system to induce and analyze tumor-specific T cell responses. Naive T cell receptor (TCR) transgenic OT-1 CD8+ T cells, which recognize the chicken ovalbumin (OVA)_257–264_ peptide, were activated for 72 hours using anti-CD3, anti-CD28, and IL-2. These activated OT-1 cells were then co-cultured with E0771 breast cancer or TC1 lung adenocarcinoma cells constitutively expressing the OVA protein, either in complete medium or in medium deprived of serine and glycine.

Co-culture in serine/glycine-deprived conditions ((-) SG) significantly reduced CD8+ T cell cytolytic activity, whereas increasing serine concentrations enhanced the response (Fig. 4c). Additionally, serine/glycine starvation decreased the expression of interferon γ (IFNγ) and granzyme B (GzmB) in OT-1 T cell co-cultures with multiple OVA-expressing cancer cell lines (Fig. 4d; Supplementary Fig. 6d-e).

These results were consistent in human cells. We co-cultured MDA-MB-231 cancer cells loaded with the melanoma antigen recognized by T cells 1 (MART-1) with T cells engineered to express a MART-1-specific T cell receptor (Supplementary Fig. 6f)^27^. In this model, serine/glycine deprivation significantly reduced tumor-reactive T cell cytotoxic activity and the expression of key T cell effector cytokines, including IL-2, TNFα, and IFNγ (Supplementary Fig. 6g-h).

Next, to investigate the dynamics of serine metabolism in co-cultures of cancer and T-cells during antigen-specific responses, we employed a cell-size filtration-based technique enabling the rapid separation of both populations (Supplementary Fig. 6c)^28^. We treated mono- and co-cultures of OT-1 CD8+ T cells and E0771-OVA cells with U-[^13^C]-glucose, and quantified the incorporation of ^13^C-glucose-derived carbon into serine and glycine using liquid chromatography-mass spectrometry (LC-MS). Intracellular pools of serine and glycine in naïve T cells incorporated a minimal fraction of glucose into serine and glycine; however, approximately 30% of the intracellular pools of serine and glycine were labelled from glucose upon antigen-specific activation (Fig. 4e-f). Conversely, glucose-dependent labelling of serine and glycine in E0771-OVA cells slightly decreased upon heterotypic interaction with OT-1 CD8^+^ T cells (Fig. 4e-f), suggesting that upon antigen recognition, T cells activated serine and glycine synthesis from glucose. These findings were consistent with the increased expression of genes associated with the serine biosynthesis pathway (PHGDH, PSAT1, and PSPH) and genes enabling serine entry into one-carbon metabolism (SHMT1 and SHMT2), which were observed exclusively in OT-I CD8+ T cells after physical interaction with OVA-expressing cancer cells or SIINFEKL-pulsed antigen presenting cells (Fig. 4g-h; Supplementary Fig. 6i-l).

Consistent with these findings, global rates of protein synthesis increased in OT-1 T cells following antigen recognition in a serine/glycine-dependent manner, while protein synthesis rates in E0771-OVA cells remained unchanged (Fig. 4i). Moreover, when measuring tRNA aminoacylation levels in co-cultures of cancer and T cell populations during antigen-specific responses (Supplementary Fig. 6c), we observed a significant decrease in the aminoacylation levels of serine (Ser)-tRNAs and glycine (Gly)-tRNAs in OT-1 T cells, whereas the levels remained unaffected in E0771-OVA cancer cells (Fig. 4j). These results were replicated in human MDA-MB-231 and MART-1 T cell co-cultures (Supplementary Fig. 6m-n). Collectively, our data demonstrate that serine and glycine are crucial for T cell protein synthesis in the TME within the context of immune checkpoint blockade and are required amino acids for an efficient T cell cytotoxic response.

### The availability of serine and glycine is a critical requirement for effective immune checkpoint blockade

Our *in vivo* DualRP datasets revealed an increase in the expression of genes linked to the serine biosynthesis pathway (PHGDH, PSAT1, and PSPH) and those enabling serine entry into one-carbon metabolism (SHMT1 and SHMT2) in the T cell compartment after the administration of αPD-1. By contrast, cancer cells did not exhibited changes (Fig. 5a), suggesting that the serine synthesis and the one carbon metabolism pathways are enhanced exclusively in the T cell compartment of tumors in response to checkpoint inhibition.

**Figure 5.**
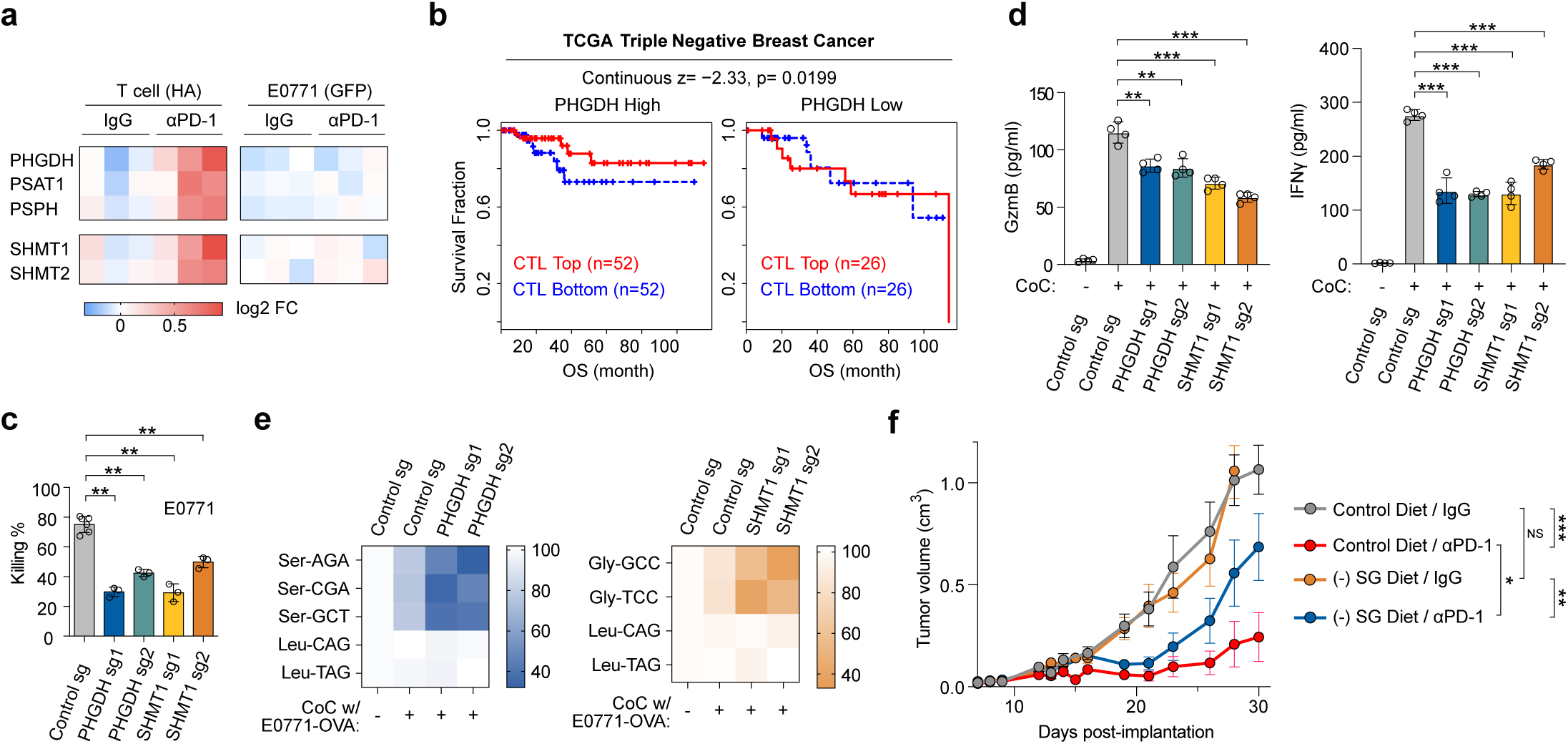
Serine and glycine are crucial for effective immunotherapy, with the serine synthesis pathway associated with T cell dysfunction in cancer patients. **a,** Heatmap depicting the log2 fold-change (FC) values of differentially expressed genes in T cells and E0771 cancer cells from tumors treated wither with IgG (2 µg/µl) or αPD-1 (2 µg/µl). **b,** TIDE analyses were conducted on PHGDH expression within T cell dysfunction signatures associated with improved survival in patients with Triple Negative Breast Cancer. **c,** Cytotoxicity assay by CD8+ T cells expressing sgRNAs targeting Phgdh or Shmt1 and co-cultured with GP_33–41_-pulsed E0771 cells for 24 hrs. Data represent mean ± SD (n = 3); **P < 0.01 by Student’s t-test. **d,** GzmB and IFNγ secretion by CD8+ T cells transduced with sgRNAs against Phgdh or Shmt1 and co-cultured with GP_33–41_-pulsed E0771 cells for 24 hrs. Data represent mean ± SD (n = 4); **P < 0.01; ***P < 0.001by Student’s t-test. **e,** tRNA aminoacylation assay in T cells expressing sgRNAs targeting Phgdh or Shmt1 and co-cultured with GP_33–41_-pulsed E0771 cells. Cells were separated by size (as described in Supplementary Fig. 6c) after 24 hours of co-culture. Ser-tRNAs, Gly-tRNAs and control Leu-tRNAs were analyzed. **f,** Mice injected orthotopically with E0771 cells were fed a control or (-) SG diet 7 days after tumor cell injection. When tumors became palpable, half of the animals in each group were treated with either IgG (2 µg/µl) or anti-PD1 (2 µg/µl), and tumor volume was monitored over time. Tumor volumes are presented as the mean ± SD (n=7); NS, non-significant; *p < 0.05; **p < 0.01; ***p < 0.001 determined through a Student’s t-test.

Next, to assess the clinical relevance of serine metabolism in cancer, we employed Tumor Immune Dysfunction and Exclusion (TIDE) analysis to investigate whether PHGDH expression in T cells correlates with outcomes across different cancer types^29^. Intriguingly, while a high cytotoxic T lymphocyte (CTL) score was associated with an overall survival benefit, low levels of PHGDH expression abolished this benefit in cases of TNBC, melanoma, lung adenocarcinoma, and endometrial tumors (Fig. 5b; Supplementary Fig. 8a-d), highlighting the importance of de novo serine synthesis in predicting cancer immunotherapy outcomes in patients.

Subsequently, we aimed to determine the essentiality of the serine synthesis and the one carbon metabolism pathways to ensure an effective cytotoxic T-cell response. To this end, we transduced Cas9-expressing CD8^+^ T cells derived from P14 x CD4-Cre x Rosa-LSL-Cas9-GFP mice with sgRNAs targeting either PHGDH or SHMT1 (Supplementary Fig. 7a-b)^30^. These T cells express a lymphocytic choreomeningitis virus (LCMV) glycoprotein 33 (GP33)-specific T cell receptor (TCR). Remarkably, we observed that T cells lacking PHGDH or SHMT1 displayed a significant reduction in cytotoxic activity when co-cultured with GP_33–41_ peptide-pulsed E0771, B16, or TC1 cells, in comparison to the control vector (Fig. 5c; Supplementary Fig. 7c). Furthermore, we studied the immunological traits of PHGDH and SHMT1 KO CD8^+^ T cells. Interestingly, these mutants exhibited decreased levels of granzyme B and IFNγ when co-cultured with pulsed E0771 cells (Fig. 5d). Moreover, tRNA aminoacylation levels of Ser- and Gly-tRNAs in CD8^+^ T cells expressing PHGDH sgRNAs were significantly reduced upon antigen recognition (Fig. 5e). Our findings collectively demonstrate that genetic perturbations in the serine synthesis and one-carbon metabolism pathways lead to diminished cytotoxic activity in CD8^+^ T cells and a reduction in the availability of serine and glycine for protein synthesis.

To investigate the significance of serine and glycine in the effective response to checkpoint blockade immunotherapy, we employed a syngeneic breast cancer model with the murine cell line E0771. After injecting cancer cells into the mammary fad pads, we transitioned the mice to either a control diet or a serine/glycine-free ((-) SG) diet. When the tumors became visible, mice were treated with either αPD-1 or IgG. Notably, dietary intervention alone did not affect tumor growth (Fig. 5f). However, administering αPD-1 as monotherapy substantially reduced the tumor burden and markedly improved survival (Supplementary Fig. 8e). When combining the (-) SG diet with αPD-1 treatment, it considerably attenuated the effects of immune checkpoint blockade (Fig. 5f; Supplementary Fig. 8e). This reduction in efficacy correlated with changes in tumor-specific T-cell numbers within the tumors (Supplementary Fig. 8f), suggesting that systemic serine and glycine are key to an efficient immunotherapeutic response. In summary, these data demonstrate the utility of DualRP not only in concurrently analyzing gene expression in two interacting cell populations *in vitro*, but also in revealing the metabolic constraints within distinct cellular compartments of tumors. This tool has enabled us to establish that serine and glycine are crucial for effective immunotherapy and that the serine synthesis pathway is associated with T cell dysfunction-related clinical outcomes in cancer patients.

## Discussion

Various cell types, including cancer-associated fibroblasts, endothelial cells, and immune cells, have a substantial impact on shaping the metabolic behavior of cancer cells^31–35^. Notably, certain TNBC cells, when in contact with stromal fibroblasts, exhibit increased expression of ISGs^36–39^, this phenotype correlated with the ability of cancer cells to acquire resistance to ionizing radiation^40,41^. Our study, utilizing the DualRP approach, provides evidence that the activation of type I IFN signaling during these heterotypic interactions results in remodeling of lysosomal function, enabling cancer cells to generate the necessary nutrients required for tRNA aminoacylation. In line with our findings, a recent study showed that type I IFN signaling enhances lysosomal function during bacterial infections^42^.

A deficiency in essential nutrients within the TME can impede T cell activation, reducing their capacity to target and eliminate tumors. For instance, limitations in amino acids such as serine, arginine, asparagine, and alanine impair T cell activation *in vitro*.^26,43–45^. Tumors with a high abundance of metabolically exhausted T cells tend to be more resistant to immune checkpoint inhibitor treatments^46–48^. Therefore, the identification and restoration of metabolic constraints within the TME have the potential to enhance the efficacy and long-term success of immunotherapy. Our study utilized the DualRP system to simultaneously investigate the metabolic constraints experienced by T cells and cancer cells during immune checkpoint blockade. We found that the T cell compartment primarily encounters limitations in serine and glycine amino acids, and we demonstrated that restricting these amino acids adversely affects T cell fitness and the antitumor immune response.

Our data suggests that upon antigen recognition, T cells activate serine and glycine synthesis from glucose. Interestingly, the labeled serine detected in T cells may originate from extracellular sources, such as export by cancer cells, rather than exclusively from glucose-derived synthesis. This alternate perspective broadens our understanding of the metabolic interplay within the TME. Our findings underscore the critical role of serine and glycine metabolism in supporting T cell fitness during immune checkpoint blockade therapy, regardless of their precise source.

Notably, we observed that all six serine codons are affected upon immune checkpoint blockade therapy. In contrast, Banh et al. reported a codon-specific reduction in mRNA translation, particularly at TCC and TCT serine codons. This discrepancy likely reflects differences in experimental conditions, such as the duration of serine deprivation and the cell types analyzed. Codon-specific effects during serine depletion have significant implications, suggesting distinct translational and metabolic responses that depend on the cellular context and nutrient availability.

CTLA-4 and PD-1 share functional similarities in suppressing T cell activity; however, their molecular mechanisms differ significantly^49^. CTLA-4 primarily acts during the early stages of T cell activation by competing with CD28 for co-stimulatory signals. This competition is crucial as CTLA-4’s higher affinity for CD80/CD86 effectively limits CD28-mediated signaling, thereby increasing the activation threshold of T cells and reducing immune responses to weak antigens such as self- and tumor antigens. In contrast, PD-1 exerts its effects later in the immune response by dampening T cell responses within the tumor microenvironment through distinct pathways that inhibit signaling cascades such as PI3K/AKT^49,50^.

Metabolically, both CTLA-4 and PD-1 have been associated with reduced glycolysis and lipid metabolism in T cells^51,52^. However, their specific effects on amino acid pathways—including serine synthesis—remain less explored. Depletion of serine leads to diminished nucleotide production, resulting in a decline in cellular proliferation^17,53^. Recent studies demonstrated that formate or methanol supplementation can enhance effector T cell fitness and substantially improve the efficiency of immunotherapy^54,55^. Our findings underscore the critical role of serine restriction not only in nucleotide production but also in tRNA charging. Importantly, it is possible to rescue tRNA charging without fully restoring protein synthesis, as protein production can still be reduced indirectly by signaling pathways that respond to changing amino acid levels, such as GCN2^56^. This suggests that ribosome stalling at specific codons—such as those for alanine or glycine—might not directly impair protein synthesis. Instead, these changes could influence signaling pathways that regulate translational processes.

Combinatorial treatments involving checkpoint inhibitors and dietary restoration of one carbon units and serine/glycine to sustain nucleotide and protein synthesis, respectively, hold great potential to enhance immune-based treatments. Recent advances have enabled the exploration of metabolic states within specific cell types, expanding our understanding of the metabolic interplay within complex cell populations^57–59^. In this study, we describe DualRP, a versatile tool that allows for the simultaneous investigation of metabolic constraints in distinct cell populations within the TME.

While our current analyses focus on amino acid limitations, DualRP inherently captures ribosome footprints across mRNAs, enabling identification of translationally regulated genes. Future studies could leverage these capabilities to systematically identify mRNAs that exhibit differential ribosome occupancy independent of transcriptional changes. For example, ribosome occupancy could be correlated with protein expression using proteomics or validated via western blot analyses. We envision that our approach can be extended to encompass multiple cellular populations, thereby facilitating the study of multidimensional metabolic interactions in cancer and organismal development.

## Methods

### Cell culture

Human cell lines (SUM-159PT, MRC5, MDA-MB-231) were cultured in DMEM High Glucose Medium (Thermo Fisher Scientific; cat. 41966-029) supplemented with 10% FBS (Thermo Fisher Scientific; cat. 10270-106) and 100 units/ml penicillin and 100 mg/ml streptomycin (Thermo Fisher Scientific; cat. 15140-122). For glucose starvation experiments, cells were cultured for 48h in DMEM without glucose and pyruvate (Thermo Fisher Scientific; cat. 11966-025) supplemented with either 4.5 g/l (high glucose) or 0.45 g/l glucose (low glucose), 10 % dialysed fetal bovine serum (Thermo Fisher Scientific; cat. 2093867) and 100 units/ml penicillin and 100 mg/ml streptomycin.

Mouse cell lines (B16, E0771 and TC1) were grown in RPM1 1640 (Thermo Fisher Scientific; cat. 21875-034) supplemented with 10 % FBS, 100 units/ml penicillin and 100 mg/ml streptomycin (Thermo Fisher Scientific; cat. 15140-122), 1 mM sodium pyruvate (Thermo Fisher Scientific; cat. 11360-070), 20 mM HEPES (Sigma; cat.H4034) and 50 µM β-mercaptoethanol (Sigma; cat. M6250). Mouse cell lines expressing chicken ovalbumin (B16-OVA and TC1-OVA) or the lymphocytic choriomeningitis virus (LCMV) glycoprotein 33 (B16-GP33) were kindly donated by Dr. Guoliang Cui and Dr. Chong Sun. Platinum-E cells were used at low passage and cultured in DMEM (Thermo Fisher Scientific; cat. 41966-029) supplemented with 10 % FBS (Thermo Fisher Scientific; cat. 10270-106), 100 units/ml penicillin and 100 mg/ml streptomycin (Thermo Fisher Scientific; cat. 15140-122) and 1% GlutaMAX (Thermo Fisher Scientific; cat. 35050061). All cell lines were regularly tested for Mycoplasma contamination and kept at 37 °C in 5 % CO2.

### Homozygous knock-in of ribosomal fluorescent proteins

Endogenous fluorescent tagging of ribosomal proteins RPL10a and RPS3 was performed in triple-negative breast cancer lines via homozygous knock-in with paired CRISPR-Cas9 as previously described^60^. In brief, two plasmids expressing the Cas9D10A nickase (Backbone Addgene: 42335) and sgRNAs targeting the ribosomal protein locus, were transfected in the presence of the DNA repair template of a donor plasmid containing the fluorescent marker flanked by homology arms (Sequences are described in Table S1). Selection of single-cell clones expressing fluorophore was carried out by FACS on a 96 well plate. Positive homozygous clones were genotyped and tested for polysome incorporation of the endogenously expressed chimera. Clonal proliferation was compared to the parental control for 8 days using the IncuCyte S3 Live-Cell Analysis System.

### Lentiviral overexpression and knock out generation

MRC5 fibroblasts expressing heterozygous ribosomal tagging (MRC5-mCherry-RPL10a) were produced via lentiviral overexpression. Virus was generated in HEK293Tx using PEI max transfection reagent (Polyscience; cat. 24765-1), lentiviral packaging plasmids (Addgene: 12251, 12253 and 14888) and the vector coding for the fluorescent chimera (pCDH1-mCherry-RPL10a). After viral transduction in presence of polybrene (Santa Cruz;cat. Sc-255611), cells were sorted for mCherry via FACS. sgRNA against STAT1 were cloned into a lentiviral expression vector for Cas9 (plentiCRISPR v2; addgene 52961). SUM159PT and MRC5 knock outs were prepared via lentiviral transduction and selected with 2 μg/ml puromycin (Sigma; cat. P8833) for 5 days. E0771 cells expressing chicken ovalbumin (E0771-OVA) were generated via lentiviral transduction as described above and selected with 10 μg/ml blasticidin (Santa Cruz; cat.sc-495389) for 3 days.

### Immunoblotting

Cell extracts were separated on 12% SDS–PAGE gels and transferred to 0.45 µm nitrocellulose membranes (Biorad; cat. 1620115). Antibodies used were RPL10a (Abcam; cat. Ab174318, 1:1,000), RPS3 (Cell Signaling; cat. 9538, 1:1,000), eGFP (Santa Cruz; cat. sc-9996, 1:1,000), mCherry (Abcam; cat. Ab167453 1:1,000), Neongreen (Chromotek; cat. 32F6, 1:1,000), RPL22 (Santa Cruz; cat. sc-373993 1:500), HA-tag (Biolegend; cat. 901501, 1:1000), STAT1 (Cell Signaling; cat. 9172, 1:1,000), PHGDH (Sigma; cat. HPA021241, 1:1,000), SHMT1 (Cell Signaling;cat. 80715, 1:1,000), LYSET (Atlas Antibodies; cat. HPA048559, 1:1,000), Cathapsin B (R&D Systems; cat. AF953, 1:1000), Cathapsin L (R&D Systems; cat. AF952 1:1000), HEXA (R&D Systems; cat. AF6237, 1:1000).

For immunoblotting of polysome fractions, cells were washed with ice-cold PBS with 100 µg/ml cycloheximide (CHX) and lysed in NP40 lysis buffer (20 mM Tris-HCl pH. 7.5 pH, 10 mM MgCl2 ,150 mM KCl, 1 % NP40, 2 mM DTT, 1x EDTA-free complete protease inhibitors and 100 µg/ml CHX). Cell extract was pipetted on top of a sucrose gradient and centrifuged for 2 hours at 36.000 rpm and 4°C in the Optima XPN 100 ultracentrifuge with swinging the bucket rotor SW41 Ti (Beckman Coulter). Fractions were loaded in 12% SDS–PAGE gels and transferred as described above.

### Dual Ribosome Profiling

SUM159-GFP-RPL10a and MRC5-mCherry-RPL10a cells were plated one million each in 15 cm dishes, and cultured for 48h. On the day of harvesting, cells were washed with ice-cold PBS supplemented with 100 μg/ml CHX, scraped, and centrifuged at 1300 x g for 5 min. Cells were lysed in 1 ml of NP40 lysis buffer (20 mM Tris-HCl, pH 7.8, 10 mM MgCl2, 150 mM KCl, 1% NP40, 2 mM DTT, 1× Complete protease Inhibitors, 100 ug/ml CHX) for 15 min on ice. Tumor tissue was snap frozen in liquid nitrogen upon surgical removal and lysed in a tissue homogenizer for 30 s at 2500 rpm (Mikro dismembrator-S), in ice-cold NP40 lysis buffer. Cell extracts were centrifuged at 4°C for 10 min at 1300 x g and the supernatant subjected to RNAse I (1 U/μl; Ambion) digestion for 30 min at room temperature under constant rotation. Resulting monosomes were purified with 40 ul GFP-Trap-, RFP-Trap-, NeonGreen-Trap-Magnetic beads (Chromotek) or protein A/G beads (Thermo Fisher Scientific) pre-loaded with HA epitope (Biolegend; cat. 901501). After 1 h incubation at room temperature, beads were washed three times with 1 ml of NP40 Lysis buffer and three times with NP40 wash buffer (20 mM Tris-HCl pH 7.8, 10 mM MgCl2, 350 mM KCl, 1% NP40, 2 mM DTT, 1× Complete protease Inhibitors, 100 μg/ml CHX). To cleave immunoprecipitated ribosomes, beads were incubated for 1 h at 45 °C in 300 μl of NP40 Lysis Buffer supplemented with 1 % SDS and 5 μl of PCR-grade proteinase K (Roche; cat. 3115828001).

The supernatant was resuspended in 1 ml TriReagent (Zymo) and libraries prepared according to the previously described ribosome profiling protocol^14^. In brief, RPFs were isolated via phenol-chloroform and size-selected (20-34 nt) from a 12% Polyacrylamide-Urea denaturing gel. After RNA gel extraction with sodium acetate and ethanol precipitation, the 3’ ends of the RPFs were dephosphorylated with T4 PNK (NEB) to allow the ligation of a pre-adenylated DNA linker with T4 Rnl2(tr) K227Q (NEB). Excess of unbound linker was removed with 5’ Deadenylase and RecJf (NEB). Samples were pooled and rRNA was depleted with either mouse or human-specific rRNA biotinylated oligo cocktail (Table S1) and MyOne Streptavidin C1 DynaBeads (Invitrogen). Reverse transcription was performed with the SuperScript III RT system (Thermo Fisher Scientific), and the resulting cDNA purified on an 8 % Polyacrylamide-Urea gel. cDNA was circularized with CircLigase II (Lucigen) following the manufacturer’s instructions. Circularized product was PCR amplified with Q5 High Fidelity Master Mix (NEB). After DNA cleaning (Zymo) and purification in a 8 % PAA-gel, library DNA concentrations were determined by Qubit HS DNA kit and adjusted to 2nM before sequencing.

### tRNA aminoacylation assay

Cells were harvested and resuspended in a solution of 0.3 M sodium acetate/acetic acid (NaOAc/HOAc) (pH 4.5) (Thermo Fisher Scientific). Total RNA was isolated using acetate-saturated phenol/CHCl3 (pH 4.5) (Thermo Fisher Scientific). RNA was resuspended in 10 mM NaOAc/HOAc (pH 4.5). Samples were split in two, one half (2 μg) was oxidized with 50 mM NaIO4 in 100 mM NaOAc/HOAc (pH 4.5) (Santa Cruz Biotechnology) for 15 min and the other half (2 μg) was incubated in 50 mM NaCl in 100 mM NaOAc/HOAc (pH 4.5) for 15 min. Samples were quenched with glucose 100 mM for 5 min at room temperature, purified in G50 columns (GE Healthcare), and then precipitated with ethanol. tRNAs were deacylated in 50 mM Tris-HCl (pH 9) for 30 min at 37 °C. RNA was precipitated and then ligated to the 3′ adaptor tRNA using T4 RNA ligase 2 (NEB) for 2 h at 37°C. Reverse transcription was performed with the high processivity SuperScript IV synthesis kit (Thermo Fisher Scientific) at 60°C for 15 minutes. To confirm that no stalling or premature termination occurred, full length tRNA sequences were validated by Sanger sequencing. Relative aminoacylation levels were calculated by qRT–PCR using tRNA specific primers (Table S1).

### tRNA-Ribosome association assay

Cells were washed with ice-cold PBS supplemented with 100 μg/ml of CHX and lysed using NP40 lysis buffer. Ribosomes were immunoprecipitated with GFP-Trap, RFP-Trap, or NeonGreen-Trap Magnetic beads (Chromotek), as previously described. RNA was isolated using saturated phenol/CHCl3 (pH 4.5) (Thermo Fisher Scientific). 500 ng of precipitated RNA was deacylated in 50 mM Tris-HCl (pH 9) for 30 minutes at 37°C. The RNA was then precipitated and ligated to the 3′ adaptor tRNA using T4 RNA ligase 2 (NEB) for 2 hours at 37°C. Reverse transcription and sequence verification were performed under the same conditions as the tRNA aminoacylation assay. Primers and linker sequences are provided in Table S1.

### Illumina sequencing

Ribo-Seq libraries were sequenced in the NextSeq2000 (Illumina) using the 100 cycles cartridge and P2 flow cell, at the DKFZ Sequencing Open Lab. According to the Illumina Systems Guide, library was diluted and loaded at a concentration of 650 pM by diluting 7.8 μl of the 2 nM library with 16.2 μl of supplied RSB. To monitor the quality of the sequencing run, 1 μl of 1 nM PhiX (Illumina) was added to the mixture, resulting in a ∼2% PhiX spike-in. The runs were set up for single-read and 110 cycles.

### Sequencing data analysis

For data preprocessing and alignment, the adaptor sequences were trimmed using cutadapt (v3.4) and demultiplexed with barcode splitter from FASTX-toolkit (Cold Spring Harbor Laboratory, Hannon Lab, by Assaf Gordon). rRNA and tRNA sequences were filtered by alignment to indices of rRNA and tRNA sequences respectively, using BLAST-Like Alignment Tool (BLAT). A rRNA index was constructed from GENCODE v19 annotations, transcript types “rRNA”, “Mt_rRNA” and “rRNA_pseudogene”, supplemented with UCSC repeats of class “rRNA”. The tRNA index was constructed from sequences obtained from GtRNAdb25 at June 2023. Unique molecular identifiers (UMIs) were extracted with umi_tools (v1.1.1) and the rRNA discarded with BLAST-Like Alignment Tool (BLAT, v36×2). The remaining reads were aligned to the GRCh37/hg19 human genome or to GRCm38/mm10 mouse genome by Spliced Transcripts Alignment to a Reference (STAR, v2.5.3a). PCR duplicates were removed for differential gene expression analysis using umi tools. The data was subjected to differential expression analysis with DESeq2 (v1.8.2) and transcripts sorted by fold change with a Benjamini-Hochberg adjusted p-value of ≤ 0.05. Gene Ontology enrichment (GO term) analysis was conducted with clusterProfiler (v3.14.3) and R (v3.6.2).

Subsequence shift analysis involves comparing the frequencies of codon occupancy by ribosome-protected fragments (RPFs) between different samples. This comparison is done at the gene level, normalizing for gene expression discrepancies to ensure that observed differences in codon frequencies are not due to variations in gene expression levels. Specifically, RPFs (>26 nt) were assigned gene IDs and reading frames using GENCODE v19/BASIC. RPFs falling outside valid coding sequences (considering the 15-nucleotide 5′-overhang), those with ambiguous gene IDs or reading frames, were excluded. The remaining RPFs were utilized to tally the frequencies of all codons at different positions (12 and 15 nucleotides from the 5′-end) across all genes in the transcriptome. Gene-specific frequencies were determined by dividing the observed counts by the total counts for each gene. These normalized codon frequencies were then averaged across genes with a minimum count threshold in both condition and control samples (set at 100 for all figures unless stated otherwise). Shifts in codon frequencies were calculated based on these normalized and averaged values, expressed as (condition – control)/control, providing a relative measure of codon shifts compared to the control. To assess significant differences in codon occupancy at the amino acid level, subsequence shifts between control and condition samples across replicates were examined. This analysis utilized a linear mixed model (implemented via the R package ’lme4’) with fixed effects for the 20 amino acids and random effects for codons, yielding t-values and P-values. Multiple testing correction was performed using the Benjamini– Hochberg method (’p.adjust’ function in R with method set to “fdr”) to obtain adjusted P-values.

For RPF density analysis, codon-regions spanning 61 nucleotides around specified codons throughout the transcriptome (as annotated by GENmethod v19/BASIC) were identified using transcript coordinates, ensuring that codon-regions were entirely contained within exons. Overlapping transcript annotations were accounted for, retaining overlapping regions while collapsing those with identical genomic coordinates. Codon-regions unable to extend to the full 61 nucleotides (typically near transcript ends) were excluded from further analysis. The 5′ ends of ribosome-protected fragments (RPFs) were counted within each codon-region. For comparative analysis between two samples, only codon-regions with a minimum count of n in both samples were considered (set at 50 for most figures, unless specified otherwise; occasionally lowered to include at least 1,000 windows for common codons). Normalized 5′ end RPF density for each sample and codon-region was computed by dividing the total counts within the region by the region’s width, ensuring an average density of 1 within each region. These normalized densities underwent convolution using a rectangular window of width 3 and height 1/3. The average density across codon-regions was calculated, and the disparity in mean densities between samples yielded the density shift.

The code for the diricore pipeline can be found at https://github.com/A-X-Smitt/B250_diricore

### Out-of-frame analysis

To evaluate the statistical significance of codon-specific shifts. Briefly, we generated a background distribution by shifting ±1 nucleotide relative to the codon under analysis. To minimize the influence of true signals, specific values were excluded from this background distribution. For instance, under glucose starvation, which produces a signal at the GCA codon in the 15th position, we excluded NGC and CAN from the 11th and 13th positions, respectively, while including other nucleotide triplets observed at these positions. This refined background distribution was then used to determine the significance of the diricore signal at the original (0) position using Z-tests. Additionally, we verified that the background values approximated a normal distribution through visual inspection of Q-Q plots, as well as the Anderson–Darling and Shapiro–Wilk tests.

### Fluorescence microscopy and image analysis

Cells were plated on 8-well chambered coverslips (IBIDI) and left to attach overnight. Lysosomal proteolysis was investigated by live cell imaging with DQ BSA green or red (Invitrogen). In brief, cells were incubated with 0.1 mg/ml DQ BSA for 4–6 h, washed two times and chased for 3 h in fresh media to allow lysosomal accumulation of DQ BSA. 0.5 µg/ml Hoechst were added prior imaging. Imaging was performed in a humidified chamber at 37 °C and 5% CO2 with a Leica TCS SP5 confocal microscope using a ×40 or ×63, 1.40 oil objective. Fluorescence was quantified using the particle analyzer function of Fiji in randomly chosen fields of view across the entirety of each sample. Mean cellular fluorescence was determined by normalizing the integrated signal density of the respective fluorescent probe to cell number. To quantify lysosomal DQ BSA fluorescence dequenching, DQ BSA integrated density was normalized to the total cell area.

For PCA, RPFs were mapped to the coding sequence (CDS). Counts were log transformed and normalized with DESeq2 (v1.8.2). PCA was performed with the prcomp function of Stats package (v.4.3.3).

### RNA isolation and quantitative real-time PCR

Total RNA isolation was performed via phenol-chloroform extraction using TRI Reagent (Zymo Reseach; cat. R2050-1-200), followed by ice-cold isopropanol precipitation and centrifugation at 20,000 x g for 45 min at 4°C. RNA pellet was washed twice with 75 % ethanol, and resuspended in nuclease-free water. The concentration was determined at 280 nm with NanoDrop One C (Thermo). 600 ng of total RNA was reversed transcribed using LunaScript RT Supermix (NEB; cat. M3010L) and quantitative real-time PCR was performed using Luna Universal qPCR Mix (NEB; cat. M3003X) following the manufacturer’s instructions. Ct values were obtained with the Quantstudio 5RT qPCR System and analyzed using Quantstudio Design and Analysis Software 2.6.0. mRNA fold change of target genes was calculated by the ΔΔCt method. mRNA expression was normalized to GAPDH in human samples and β-Actin in mouse samples.

### Flow cytometry and flow sorting

Flow cytometry was performed on the BD Canto or Fortessa HTS FACS machines. FACS sorting was conducted on a BD Aria II and flow analyses were performed with FlowJo 10.6.1 software (LLC). For rapid separation of CD8+ T cells and tumor cells from co-culture plates, CD8 T cells were transferred from the co-culture plates to a 10 μm cell strainer (PluriSelect) to retain any tumor cells detached from the plate as previously described^28^. Staining with 1:100 APC anti-mouse CD8, clone 53-6.7 (Invitrogen) and 1:1000 DAPI (Thermo Fisher Scientific) was carried out for 1 h at 37°C and living cells were sorted with a 100 μM nozzle for GFP (cancer cells) and APC (T cells). P14:Cas9 CD8 T cells with KO of Shmt1 or Phgdh were sorted based on the expression of Cas9-GFP and expression of the BFP reporter protein coded on the sgRNA transfer vector after transduction. To quantify tumor-infiltrating lymphocytes, tumors were digested using the Tumor Dissociation Kit (Miltenyi Biotec). The cells were stained with anti-mouse CD8 antibodies (BD Biosciences, 557668, 1:200) or anti-mouse Ki-67 (BioLegend, 652424, 1:200) and analyzed using the Fortessa HTS FACS instrument.

### CD8+ T cell isolation and culture

Primary naive CD8+ T cells were purified using the MojoSort Mouse CD8 T cell isolation kit (BioLegend; cat. 480007) and activated for 72h on plates coated with 2 μg/ml anti-CD3 (BioXCell; cat. BE0001-1) and 2 μg/ml anti-CD28 (BioXCell; cat. BE0015-1) for 72 hours at 37°C. T cells were kept in RPM1 1640 (Thermo Fisher Scientific; cat. 21875-034) supplemented with 10 % FBS and 1 % penicillin-streptomycin, 1mM sodium pyruvate (Thermo Fisher Scientific; cat. 11360-070), 20 mM HEPES (Sigma; cat. H4034), 50 µM β-mercaptoethanol (Sigma; cat.M6250) and 10 ng/ml murine IL-2 (Biolegend; cat. 575404). For serine/glycine starvation experiments, medium was prepared fresh from RPMI 1640 powder without amino acids (US Biological; cat. R9010-01) and each component added individually according to the ATCC formulation. Human MART-1-specific (TCR clone #1D3) CD8+ T cells were donated by Dr. Chong Song and kept in culture as previously described^61^.

### T cell knock out generation

Platinum-E cells were transfected with pMSCV vectors (Addgene:102796) containing a BFP selection marker and sgRNAs targeting PHGDH or SHMT1. Transfection was performed using PEI max transfection reagent (Polyscience; cat. 24765-1) and 10µg DNA for 16h at 37 °C and 5% CO2. Subsequently, the cells were cultured for 8 hours in RPMI supplemented with 3mM sodium butyrate. Retroviral supernatants were collected after 48 and 72 hours transfection and filtered through 0.45µm. Naive CD8+ T cells were isolated from P14 x CD4-Cre x Rosa-LSL-Cas9-GFP mice using the MojoSort Mouse CD8 T cell isolation kit (BioLegend; cat. 480007) and activated on plates coated with 0.5 μg/ml anti-CD3 (BioXCell; cat. BE0001-1) and 2.5 μg/ml anti-CD28 (BioXCell; cat. BE0015-1) in the presence of mouse interleukins at 1 ng/ml IL-2 (Biolegend; cat. 575404), 5 ng/ml IL-7 (Peprotech; cat. 217-17) and 5 ng/ml IL-15 (Peprotech; cat. 210-15) and 55 µM β-mercaptoethanol. After 24 h activation, transduction was performed by spinfecting T cells at 2000 x g for 90 min at 30°C in presence of the retroviral supernatants and retronectin (Takara; cat. T100A). The cells were incubated at 37°C, 5% CO_2_ for 24 h before washing off the virus and expanded for 48h in medium containing 1 μg/ml anti-CD3 (BioXCell; cat. BE0001-1), 0.5 μg/ml anti-CD28 (BioXCell; cat. BE0015-1) and mouse interleukins at the concentrations mentioned above. Transduced CD8+ T cells were selected by sorting by FACS based on GFP and BFP expression.

### T cell killing and gene expression assays

CD8+ OT-I T cells were co-cultured with mouse cancer cells expressing chicken ovalbumin (B16-OVA, TC1-OVA and E0771-OVA) with increasing concentrations of serine. After 24h at 37°C and 5% CO2, cells were washed with PBS and stained with crystal violet to assess killing efficiency. Imaging was performed with Dual Lens System V850 Pro Scanner (Epson) and colony area was quantified with a previously published ImageJ plugin^62^. To determine gene expression, OT-1 T cells were co-cultured with OVA-expressing cells in a 1:4 ratio (CD8+ T cells: tumor cells) for 24h and sorted for viable CD8+ cells by FACS, pellets were subjected to RNA isolation and qPCR for serine and one carbon metabolism pathway genes. P14 T cells knock outs were co-cultured for 24h with mouse cancer cells expressing the lymphocytic choreomeningitis virus (LCMV) glycoprotein 33 (B16-GP33) or with cancer cells preloaded with LCMV gp33 peptide (IBA; cat. 6-7016-901), crystal violet staining and imaging was performed as described above. Cocultures of MART-1-specific CD8+ T cells and MDA-MB-231 MART-1 epitope transduced cells were prepared at a ratio of 1:8. After incubation for 24h and T cell removal, tumor cell viability was determined using the CellTiter Blue Assay (Promega, Cat# G8020).

### ELISA assays

Cytokine release in CD8+ T cells was quantified from cell supernatant using ELISA MAX Deluxe Set Mouse IFN-γ (BioLegend; cat. 430815) and Granzyme B Mouse ELISA (Invitrogen; cat. 88-8022-22) according to manufacturer’s instructions. Each sample was measured with the Multiskan FC (Thermo Fisher Scientific) plate reader as the 450nm-570nm absorbance’s subtraction and final concentrations were calculated with the 4-parameter logistic curve-fitting algorithm in GraphPad Prism.

### Metabolomics

CD8+ T cells were isolated from the spleen of eight-week-old OT-1 female mice and activated as previously described^28^. For the co-culture assay, 1,000,000 E0771-OVA cells/replicate were plated in 15 cm dishes 48 hours before the co-culture to let them adhere. Pre-activated OT-1 CD8+ T cells were transferred on top of the E0771-OVA cells in a 1:5 ratio (CD8+ T cells : tumor cells). For T cell-only assays, 2,000,000 pre-activated CD8+ T cells/replicate were cultured in 6-well plates. For tumor-cell-only assays, 200,000 E0771-OVA cells/replicate were plated in 6-well plates 48 hours before the assay to let them adhere. Finally, 800,000 naïve CD8+ T cells/replicate were cultured in 24-well plates. All metabolic assays were conducted in full RPMI or RPMI without serine (Bioquote, R9660). In both media, 12C glucose was replaced by 13C6 labelled glucose (Cambridge Isotope Laboratories, 110187-42-3). After six hours of culture in media with or without serine and in the presence of 13C6 glucose, CD8+ T cells and tumor cells were collected for further metabolic analysis. Collection, quenching, and metabolite extraction were performed as previously described^28^. Metabolites were measured using gas chromatography-mass spectrometry as previously described^63^.

### Analysis of protein synthesis rates

Cells were treated with as previously described^64^. Cells were treated with O-propargyl-puromycin (OPP) (20 μM, Jena Bioscience) for 1 hour at 37°C, 5% CO2. For collection, cells were washed with PBS and digested in trypsin-EDTA. For co-cultures with T cells and tumor cells, rapid separation using a 10 mm cell strainer (PluriSelect) was performed as previously described^28^. After fixing in 0.5 ml 1% w/v PFA (Sigma-Aldrich) in PBS for 15 min on ice in the dark, samples were washed in PBS and permeabilized in PBS with 3% FBS and 0.1% saponin (Sigma-Aldrich) for 5 min at room temperature. OPP was conjugated to a fluorochrome via azide-alkyne cycloaddition for 30 min at room temperature in the dark, using the Click-iT Cell Reaction Buffer Kit (Thermo Fisher Scientific and 5 μM Alexa Fluor 488/647-Azide (Thermo Fisher Scientific). Excess reagents were removed by washing cells twice in PBS with 3% FBS and 0.1% saponin. Samples were resuspended in PBS and analyzed using a BD LSRFortessa flow cytometer (BD Biosciences). Raw fluorescence median values were obtained by FlowJo 10.6.1 software (LLC) from selected subpopulations.

### Mice

6-8 weeks NSG, C57BL/6J, OT-1, Rpl22 x CD4 Cre and P14 x CD4-Cre x Rosa-LSL-Cas9-GFP were either purchased from the Charles River Laboratory or bred in our animal facility. All mice were maintained in a pathogen-free facility and used according to the German Cancer Research Center and following permission by the controlling government office (Regierungspräsidium Karlsruhe) according to the German Animal Protection Law, and in compliance with the EU Directive on animal welfare, Directive 2010/63/EU.

### Animal experiments

For tumor xenografts, orthotopic injection in the fat pad of NSG mice was performed with 2 million SUM159-eGFP-RPL10a breast cancer cells and/or in combination with MRC5-mCherry-RPL10a fibroblasts, at least 6 mice per condition were evaluated. To evaluate the effect of checkpoint inhibition in a T cell ribotag transgenic model, Rpl22 X CD4 Cre mice were injected in the fat pad with 2 million E0771-eGFP-RPL10a syngeneic breast cancer cells. Upon tumor formation, mice received two intraperitoneal injections with 3 days gap of 2 μg/μl of mouse anti-PD1 (BioXcell, BP0273) or rat IgG as control (BioXcell, BP0273), at least 4 mice per condition were evaluated. For xenografts and transgenic models, tumor growth was measured every 2-4 days until a maximum volume of 1,5 cm^3^ or 1.5 cm in one dimension and cryopreserved for DualRP library preparation. For dietary intervention experiments, C57BL/6J mice received serine-glycine free diet (TestDiet, cat. 5BQS) or full control diet (TestDiet, cat. 5BQT) after oneweek injection with the same syngeneic E0771-eGFP-RPL10a line. Mice were treated with 5 rounds of anti-PD1 or IgG with 2 days apart using concentrations above. At least 9 mice were tested per group. Experiment was ended until tumors reached a maximum volume of 1,5 cm^3^ or 1.5 cm in one dimension.

## Data availability

The sequence data from this study have been submitted to NCBI BioProject (http://www.ncbi.nlm.nih.gov/bioproject) under BioProject number PRJNA1024949.

## Acknowledgements

We thank William Faller and Joana Silva for sharing reagents. We are grateful to Aurelio Teleman for comments on the manuscript. This work was funded in part by grants of the European Research Council “DualRP” (ERC StG No. 759579) and the German Research Foundation (DFG 504774163 and DFG 545215964) to F.L.-P. D.A.H., G.P., C.P. are supported by Cancer Transitional Research and EXchange Program (Cancer-TRAX) within the German-Israeli Helmholtz International Research School. European Research Council “DRILL” (ERC StG No. 101078722) to C.S. I.E. acknowledges funding from FWO (G065122N). A.K is supported by a fellowship of the Helmholtz International Graduate School. C.C.A.R. is supported by the DKFZ International Postdoc Program.

## Author contributions

F.L.-P. conceived the project, designed all the experiments, and wrote the manuscript. Methodology and data acquisition: D.A.H., R.D.P., A.K., A.H.B., G.P., E.S., C.C.A.R., S.D., E.R., C.P., N.S., V.C.V., H.E., S.D., S.C-D., and B.M. Manuscript revision: F.L-P.

## Ethics declarations. Competing interests

The authors declare no competing financial interests.

## References

1. de Visser, K. E. & Joyce, J. A. The evolving tumor microenvironment: From cancer initiation to metastatic outgrowth. Cancer Cell 41, 374–403 (2023).

2. Dey, P., Kimmelman, A. C. & DePinho, R. A. Metabolic Codependencies in the Tumor Microenvironment. Cancer Discov. 11, 1067–1081 (2021).

3. Finicle, B. T., Jayashankar, V. & Edinger, A. L. Nutrient scavenging in cancer. Nat. Rev. Cancer 18, 619–633 (2018).

4. Reinfeld, B. I. et al. Cell-programmed nutrient partitioning in the tumour microenvironment. Nature 593, 282–288 (2021).

5. Chang, C.-H. et al. Metabolic Competition in the Tumor Microenvironment Is a Driver of Cancer Progression. Cell 162, 1229–1241 (2015).

6. Leone, R. D. & Powell, J. D. Metabolism of immune cells in cancer. Nat. Rev. Cancer 20, 516–531 (2020).

7. Reina-Campos, M., Scharping, N. E. & Goldrath, A. W. CD8+ T cell metabolism in infection and cancer. Nat. Rev. Immunol. 21, 718–738 (2021).

8. DePeaux, K. & Delgoffe, G. M. Metabolic barriers to cancer immunotherapy. Nat. Rev. Immunol. 21, 785–797 (2021).

9. Loayza-Puch, F. et al. Tumour-specific proline vulnerability uncovered by differential ribosome codon reading. Nature 530, 490–494 (2016).

10. Darnell, A. M., Subramaniam, A. R. & O’Shea, E. K. Translational Control through Differential Ribosome Pausing during Amino Acid Limitation in Mammalian Cells. Mol. Cell 71, 229–243.e11 (2018).

11. Kay, E. J. et al. Cancer-associated fibroblasts require proline synthesis by PYCR1 for the deposition of pro-tumorigenic extracellular matrix. Nat Metab 4, 693–710 (2022).

12. Banh, R. S. et al. Neurons Release Serine to Support mRNA Translation in Pancreatic Cancer. Cell 183, 1202–1218.e25 (2020).

13. Heiman, M. et al. A translational profiling approach for the molecular characterization of CNS cell types. Cell 135, 738–748 (2008).

14. McGlincy, N. J. & Ingolia, N. T. Transcriptome-wide measurement of translation by ribosome profiling. Methods 126, 112–129 (2017).

15. Dittmar, K. A., Sørensen, M. A., Elf, J., Ehrenberg, M. & Pan, T. Selective charging of tRNA isoacceptors induced by amino-acid starvation. EMBO Rep. 6, 151–157 (2005).

16. Pavlova, N. N. et al. Translation in amino-acid-poor environments is limited by tRNAGln charging. Elife 9, (2020).

17. Geeraerts, S. L., Heylen, E., De Keersmaecker, K. & Kampen, K. R. The ins and outs of serine and glycine metabolism in cancer. Nat Metab 3, 131–141 (2021).

18. Gray, L. R., Tompkins, S. C. & Taylor, E. B. Regulation of pyruvate metabolism and human disease. Cell. Mol. Life Sci. 71, 2577–2604 (2014).

19. Possemato, R. et al. Functional genomics reveal that the serine synthesis pathway is essential in breast cancer. Nature 476, 346–350 (2011).

20. Ivashkiv, L. B. & Donlin, L. T. Regulation of type I interferon responses. Nat. Rev. Immunol. 14, 36–49 (2014).

21. Reis, R. C., Sorgine, M. H. & Coelho-Sampaio, T. A novel methodology for the investigation of intracellular proteolytic processing in intact cells. Eur. J. Cell Biol. 75, 192– 197 (1998).

22. Commisso, C. et al. Macropinocytosis of protein is an amino acid supply route in Ras-transformed cells. Nature 497, 633–637 (2013).

23. Palm, W. et al. The Utilization of Extracellular Proteins as Nutrients Is Suppressed by mTORC1. Cell 162, 259–270 (2015).

24. Pechincha, C. et al. Lysosomal enzyme trafficking factor LYSET enables nutritional usage of extracellular proteins. Science 378, eabn5637 (2022).

25. Sanz, E. et al. Cell-type-specific isolation of ribosome-associated mRNA from complex tissues. Proc. Natl. Acad. Sci. U. S. A. 106, 13939–13944 (2009).

26. Ma, E. H. et al. Serine Is an Essential Metabolite for Effector T Cell Expansion. Cell Metab. 25, 482 (2017).

27. Sang, M., Lian, Y., Zhou, X. & Shan, B. MAGE-A family: attractive targets for cancer immunotherapy. Vaccine 29, 8496–8500 (2011).

28. Elia, I. et al. Tumor cells dictate anti-tumor immune responses by altering pyruvate utilization and succinate signaling in CD8+ T cells. Cell Metab. 34, 1137–1150.e6 (2022).

29. Jiang, P. et al. Signatures of T cell dysfunction and exclusion predict cancer immunotherapy response. Nat. Med. 24, 1550–1558 (2018).

30. Henriksson, J. et al. Genome-wide CRISPR Screens in T Helper Cells Reveal Pervasive Crosstalk between Activation and Differentiation. Cell 176, 882–896.e18 (2019).

31. Sousa, C. M. et al. Pancreatic stellate cells support tumour metabolism through autophagic alanine secretion. Nature 536, 479–483 (2016).

32. Vitale, I., Manic, G., Coussens, L. M., Kroemer, G. & Galluzzi, L. Macrophages and Metabolism in the Tumor Microenvironment. Cell Metab. 30, 36–50 (2019).

33. Longchamp, A. et al. Amino Acid Restriction Triggers Angiogenesis via GCN2/ATF4 Regulation of VEGF and H2S Production. Cell 173, 117–129.e14 (2018).

34. Yang, L. et al. Targeting Stromal Glutamine Synthetase in Tumors Disrupts Tumor Microenvironment-Regulated Cancer Cell Growth. Cell Metab. 24, 685–700 (2016).

35. Zhu, Z. et al. Tumour-reprogrammed stromal BCAT1 fuels branched-chain ketoacid dependency in stromal-rich PDAC tumours. Nat Metab 2, 775–792 (2020).

36. McLaughlin, M. et al. Inflammatory microenvironment remodelling by tumour cells after radiotherapy. Nat. Rev. Cancer 20, 203–217 (2020).

37. Minn, A. J. Interferons and the Immunogenic Effects of Cancer Therapy. Trends Immunol. 36, 725–737 (2015).

38. Boelens, M. C. et al. Exosome transfer from stromal to breast cancer cells regulates therapy resistance pathways. Cell 159, 499–513 (2014).

39. Nabet, B. Y. et al. Exosome RNA Unshielding Couples Stromal Activation to Pattern Recognition Receptor Signaling in Cancer. Cell 170, 352–366.e13 (2017).

40. Weichselbaum, R. R. et al. An interferon-related gene signature for DNA damage resistance is a predictive marker for chemotherapy and radiation for breast cancer. Proc. Natl. Acad. Sci. U. S. A. 105, 18490–18495 (2008).

41. Khodarev, N. N. et al. STAT1 is overexpressed in tumors selected for radioresistance and confers protection from radiation in transduced sensitive cells. Proc. Natl. Acad. Sci. U. S. A. 101, 1714–1719 (2004).

42. Zhang, H., Zoued, A., Liu, X., Sit, B. & Waldor, M. K. Type I interferon remodels lysosome function and modifies intestinal epithelial defense. Proc. Natl. Acad. Sci. U. S. A. 117, 29862–29871 (2020).

43. Geiger, R. et al. L-Arginine Modulates T Cell Metabolism and Enhances Survival and Anti-tumor Activity. Cell 167, 829–842.e13 (2016).

44. Wu, J. et al. Asparagine enhances LCK signalling to potentiate CD8+ T-cell activation and anti-tumour responses. Nat. Cell Biol. 23, 75–86 (2021).

45. Ron-Harel, N. et al. T Cell Activation Depends on Extracellular Alanine. Cell Rep. 28, 3011–3021.e4 (2019).

46. Mariathasan, S. et al. TGFβ attenuates tumour response to PD-L1 blockade by contributing to exclusion of T cells. Nature 554, 544–548 (2018).

47. Sugiura, A. & Rathmell, J. C. Metabolic Barriers to T Cell Function in Tumors. J. Immunol. 200, 400–407 (2018).

48. Hegde, P. S. & Chen, D. S. Top 10 Challenges in Cancer Immunotherapy. Immunity 52, 17–35 (2020).

49. Buchbinder, E. I. & Desai, A. CTLA-4 and PD-1 pathways: Similarities, differences, and implications of their inhibition. Am. J. Clin. Oncol. 39, 98–106 (2016).

50. Pardoll, D. M. The blockade of immune checkpoints in cancer immunotherapy. Nat. Rev. Cancer 12, 252–264 (2012).

51. Wei, S. C., Duffy, C. R. & Allison, J. P. Fundamental mechanisms of immune checkpoint blockade therapy. Cancer Discov. 8, 1069–1086 (2018).

52. Shyer, J. A., Flavell, R. A. & Bailis, W. Metabolic signaling in T cells. Cell Res. 30, 649– 659 (2020).

53. Tajan, M. et al. Serine synthesis pathway inhibition cooperates with dietary serine and glycine limitation for cancer therapy. Nat. Commun. 12, 366 (2021).

54. Rowe, J. H. et al. Formate supplementation enhances anti-tumor CD8+ T cell fitness and efficacy of PD-1 blockade. Cancer Discov. (2023) doi:10.1158/2159-8290.CD-22-1301.

55. Xu, X., et al. One-carbon unit supplementation fuels tumor-infiltrating T cells and augments checkpoint blockade. bioRxivorg (2023) doi:10.1101/2023.11.01.565193.

56. Dong, J., Qiu, H., Garcia-Barrio, M., Anderson, J. & Hinnebusch, A. G. Uncharged tRNA activates GCN2 by displacing the protein kinase moiety from a bipartite tRNA-binding domain. Mol. Cell 6, 269–279 (2000).

57. Rappez, L. et al. SpaceM reveals metabolic states of single cells. Nat. Methods 18, 799– 805 (2021).

58. Lau, A. N. et al. Dissecting cell-type-specific metabolism in pancreatic ductal adenocarcinoma. Elife 9, (2020).

59. Rodríguez-Colman, M. J. et al. Interplay between metabolic identities in the intestinal crypt supports stem cell function. Nature 543, 424–427 (2017).

60. Koch, B. et al. Generation and validation of homozygous fluorescent knock-in cells using CRISPR-Cas9 genome editing. Nat. Protoc. 13, 1465–1487 (2018).

61. Miao, B. et al. CMTM6 shapes antitumor T cell response through modulating protein expression of CD58 and PD-L1. Cancer Cell 41, 1817–1828.e9 (2023).

62. Guzmán, C., Bagga, M., Kaur, A., Westermarck, J. & Abankwa, D. ColonyArea: an ImageJ plugin to automatically quantify colony formation in clonogenic assays. PLoS One 9, e92444 (2014).

63. Rossi, M. et al. PHGDH heterogeneity potentiates cancer cell dissemination and metastasis. Nature 605, 747–753 (2022).

64. Liu, J., Xu, Y., Stoleru, D. & Salic, A. Imaging protein synthesis in cells and tissues with an alkyne analog of puromycin. Proc. Natl. Acad. Sci. U. S. A. 109, 413–418 (2012).

