## Supplementary information for "Dual Ribosome Profiling reveals metabolic limitations of cancer and stromal cells in the tumor microenvironment"

#### Supplementary Figures 1 to 8

### Supplementary Figure 1.

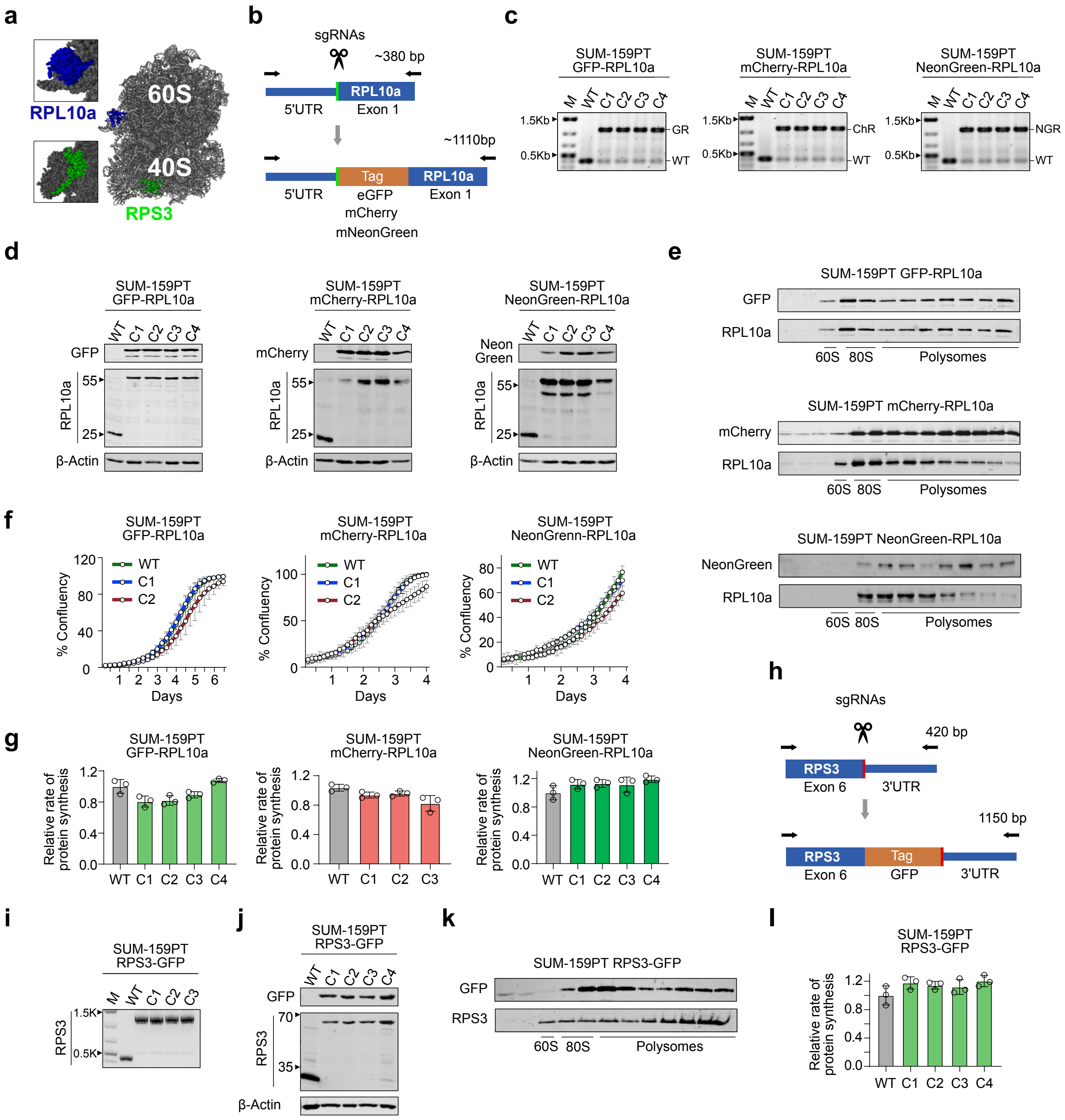

**Supplementary Figure 1. Characterization of cell lines carrying tagged ribosomes.**

**a,** The fluorescent tags indicated were endogenously incorporated at the N-terminus of *RPL10a* or the C-terminus of *RPS3*. The figure shows the location of both proteins in the large and small subunit of the ribosome, respectively.

**b,** Paired CRISPR–Cas9 nickase approach for tagging the RPL10a gene.

**c,** Gene editing was performed on SUM-159PT cells using CRISPR-Cas9 to tag either GFP-RPL10a, mCherry-RPL10a, or mNeonGreen-RPL10a to the N-terminus of RPL10a. Homozygous incorporation was confirmed through genotyping (GR, GFP-RPL10a; ChR, mCherry-RPL10a; NGR, mNeonGreen-RPL10a).

**d,** Immunoblotting was performed on lysates from clones of the indicated cells, comparing WT with tagged-RPL10a.

**e,** Western blot analysis was performed using proteins isolated from individual fractions across the sucrose gradients of cell lysates from SUM-159PT cells expressing GFP-RPL10a, mCherry-RPL10a, or NeonGreen-RPL10a.

**f,** IncuCyte cell proliferation curves of SUM-159PT clones expressing GFP-RPL10a, mCherry-RPL10a, or NeonGreen-RPL10a. Data represent mean ± SD (n = 7).

**g,** Protein synthesis rates were determined based on O-propargyl-puromycin (OP-Puro) incorporation in SUM-159PT clones expressing GFP-RPL10a, mCherry-RPL10a, or NeonGreen-RPL10a. Data represent mean ± SD (n = 3).

**h,** GFP was endogenously tagged at the C-terminus of RPS3 using a paired CRISPR–Cas9 nickase approach.

**i,** Genotyping of the homozygous incorporation of GFP in the C terminus of RPS3.

**j,** Western blot analysis of SUM-159PT WT or RPS3-GFP cell lysates.

**k,** Polysomal distribution of RPS3-GFP examined by Western blot analysis.

**l,** OP-Puro incorporation assay in SUM-159PT clones expressing RPS3-GFP. Data represent mean ± SD (n = 3).

#### Supplementary Figure 2.

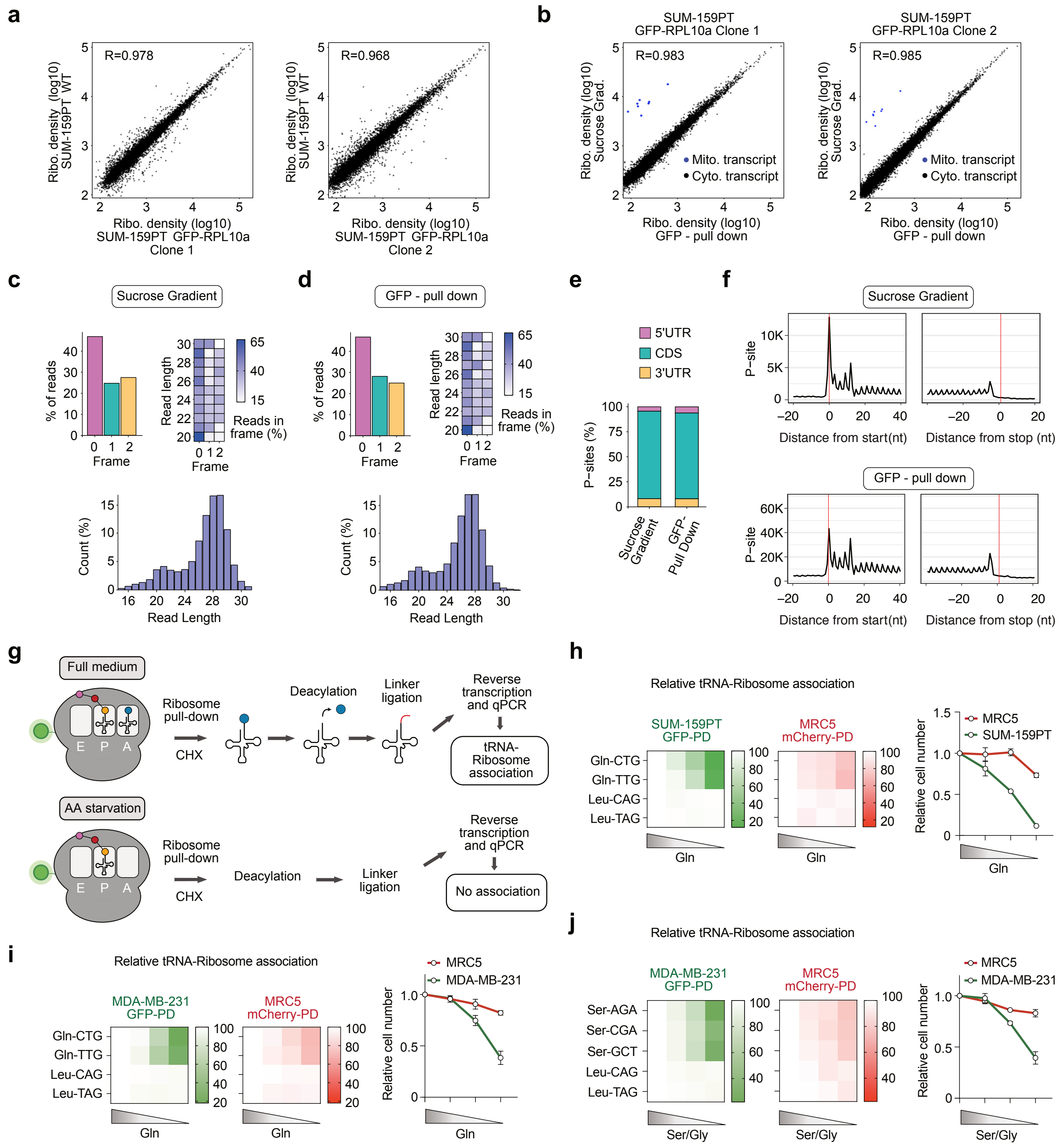

**Supplementary Figure 2. Ribosome profiling and tRNA association studies in tagged breast cancer cell lines.**

**a,** Scatter plot comparing RPF abundance between ribosome profiling datasets from SUM-159PT and SUM-159PT GFP-RPL10a clones. The Pearson correlation coefficient (R) is marked at the upper left corner. All libraries were prepared with conventional sucrose gradients.

**b,** Scatter plot comparing RPF abundance between ribosome profiling data sets generated from conventional sucrose gradients and GFP pull-downs from SUM-159PT-GFP-RPL10a clones. The Pearson correlation coefficient (R) is marked at the upper left corner. mtDNA-encoded transcripts (Mito. transcript) are underrepresented in the GFP pull-down libraries.

**c-d,** Frame and read-length distributions of the 5' end of RPFs from sucrose gradients and GFP pull-down libraries.

**e,** Percentage of reads mapping to the 5′ UTR, coding sequence (CDS), and 3′ UTR of mRNAs from data sets generated from conventional sucrose gradients or GFP pull-downs from SUM-159PT-GFP-RPL10a clones.

**f,** RPF Coverage around START and STOP codons in ribosome profiling libraries derived from sucrose gradients and GFP pull-downs.

**g,** Schematic of the tRNA-Ribosome association approach. CHX, Cycloheximide.

**h-j,** Ribosome-tRNA association assays in SUM-159PT-GFP-RPL10a (h) or MDA-MB-231-GFP-RPL10a (i-j) and MRC5-mCherry-RPL10a cells starved of glutamine (ranging from 4mM to 0mM) or serine/glycine (ranging from 0.4mM to 0mM each). The right panels display normalized cell numbers of the indicated cell lines at different concentrations of glutamine or serine/glycine. Normalized cell numbers at different serine/glycine concentrations. The data are expressed as relative cell numbers compared to the untreated group at the endpoint. Measurements were taken 48 hours after plating. PD, pull-down.

**Supplementary Figure 3.**

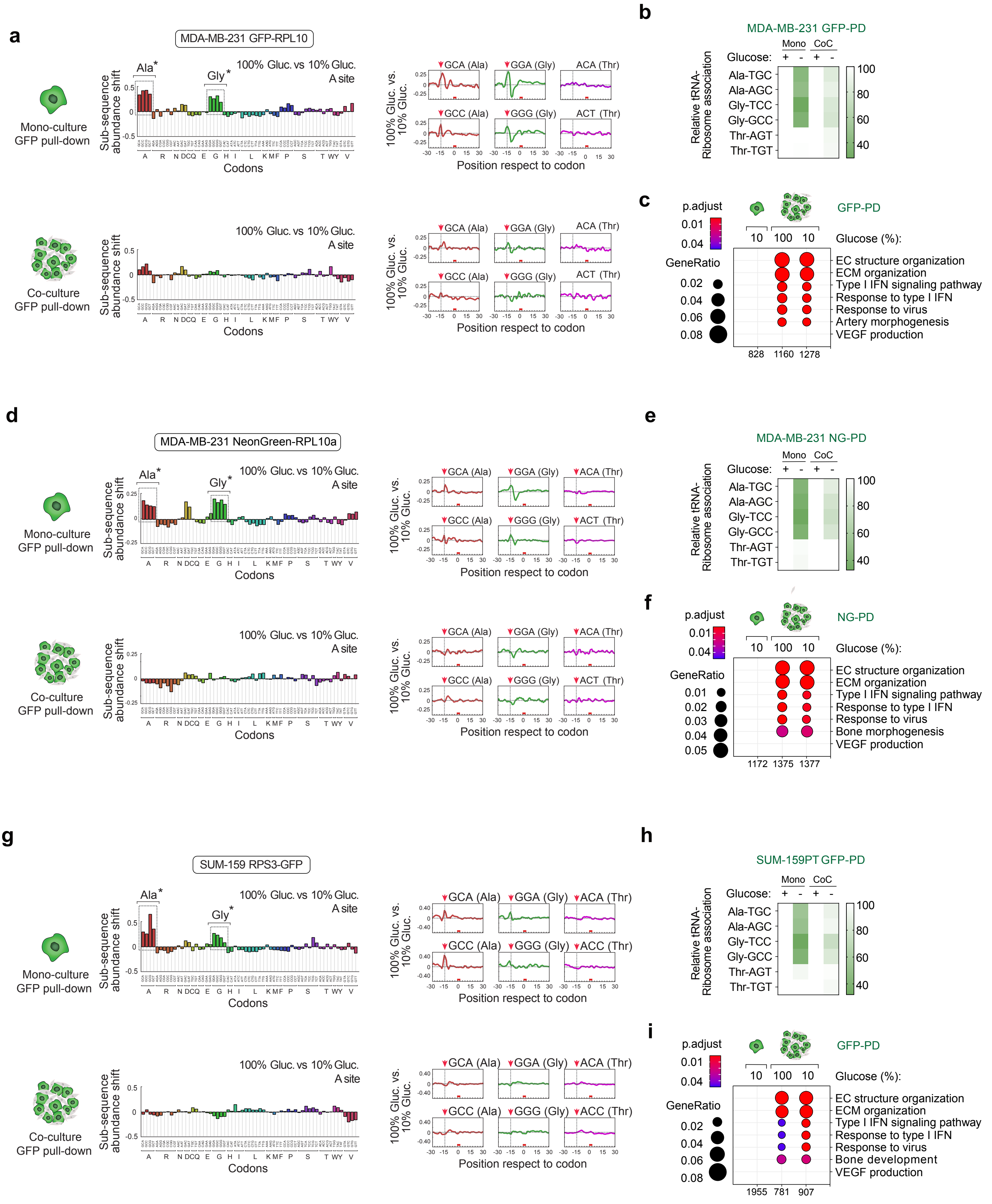

**Supplementary Figure 3. DualRP measurements are independent of cell line, protein tag, or ribosomal subunit tagging.**

**a,** Diricore analysis of GFP pull-downs from mono-cultures of MDA-MB-231-GFP-RPL10a cells (upper panel) and co-cultures with MRC5 mCherry-RPL10a (lower panel) grown in full (100%, 25mM) or glucose-deprived (10%, 2,5 mM) medium. Cells were treated for 48 hrs. The dip observed at position -12 in the ribosome density plots (right panels) reflects a reduced ribosome density at the P site of the codon. *Out-of-frame analysis P < 0.01 for the GCA, GCC, GCG, GCT Ala codons and for the GGA, GGC, GGG Gly codons.

**b,** tRNA-Ribosome association assay in mono- and co-cultures of MDA-MB-231-GFP-RPL10a and MRC5-mCherry-RPL10a growing in full ((+), 25mM) or glucose-deprived ((-), 2,5mM) medium. Data represent mean ± SD (n=3). PD, pull-down.

**c,** Gene ontology analysis of genes upregulated upon heterotypic interaction in MDA-MB-231 GFP-RPL10a grown in full (100%, 25mM) or glucose-deprived (10%, 2,5mM) medium. Significantly enriched gene sets (p-value < 0.05) are shown, along with the number of identified proteins in the respective gene set (GeneRatio). PD, pull-down.

**d,** Diricore analysis was conducted on NeonGreen pull-downs from mono- and co-cultures. The mono-cultures involved MDA-MB-231 NeonGreen-RPL10a cells (upper panel), while the co-cultures included MRC5 mCherry-RPL10a (lower panel). The cells were cultured in either full medium (100%, 25mM) or medium deprived of glucose (10%, 2.5 mM). The cells were subjected to a 48-hour treatment. *Out-of-frame analysis P < 0.01 for the GCA, GCC, GCG, GCT Ala codons and for the GGA, GGC, GGG, GGT Gly codons.

**e,** Quantification of the association of tRNA-Ribosome in both mono- and co-cultures of MDA-MB-231-NeonGreen-RPL10a and MRC5-mCherry-RPL10a cells. These cultures were maintained in two different conditions: full ((+), 25mM) medium and glucose-deprived ((-), 2.5mM) medium. Data represent mean ± SD (n=3). NG, NeonGreen; PD, pull-down.

**f,** Gene ontology analysis of genes that showed an upregulation in MDA-MB-231 NeonGreen-RPL10a cells during heterotypic interaction. This analysis was carried out under two different growth conditions: full (100%, 25mM) medium and glucose-deprived (10%, 2.5mM) medium. The results presented include significantly enriched gene sets (p-value < 0.05), accompanied by the number of proteins identified within each respective gene set (GeneRatio). NG, NeonGreen; PD, pull-down.

**g,** Diricore analysis of mono- and co-cultures of SUM-159PT RPS3-GFP with MRC5 mCherry-RPL10a cells, represented in the upper and lower panels, respectively. These cell populations were cultivated under two conditions: full (100%, 25mM) medium and glucose-deprived (10%, 2.5 mM) medium. The cells were subjected to this treatment for 48 hours. *Out-of-frame analysis P < 0.01 for the GCA, GCC, GCG, GCT Ala codons and for the GGA, GGC, GGG Gly codons.

**h,** tRNA-Ribosome association assay in mono- and co-cultures of SUM-159PT RPS3-GFP and MRC5-mCherry-RPL10a growing in full ((+), 25mM) or glucose-deprived ((-), 2,5mM) medium. Data represent mean ± SD (n=3). PD, pull-down.

**i,** Gene ontology analysis was performed on genes that exhibited an upregulation during the heterotypic interaction of SUM-159PT-RPS3-GFP with MRC5 mCherry-RPL10a cells. This analysis was conducted under two distinct growth conditions: full medium (100%, 25mM) and glucose-deprived medium (10%, 2.5mM). The results highlight gene sets that demonstrated a significant enrichment (p-value < 0.05), and the count of proteins identified within each specific gene set is presented (GeneRatio). PD, pull-down.

#### Supplementary Figure 4

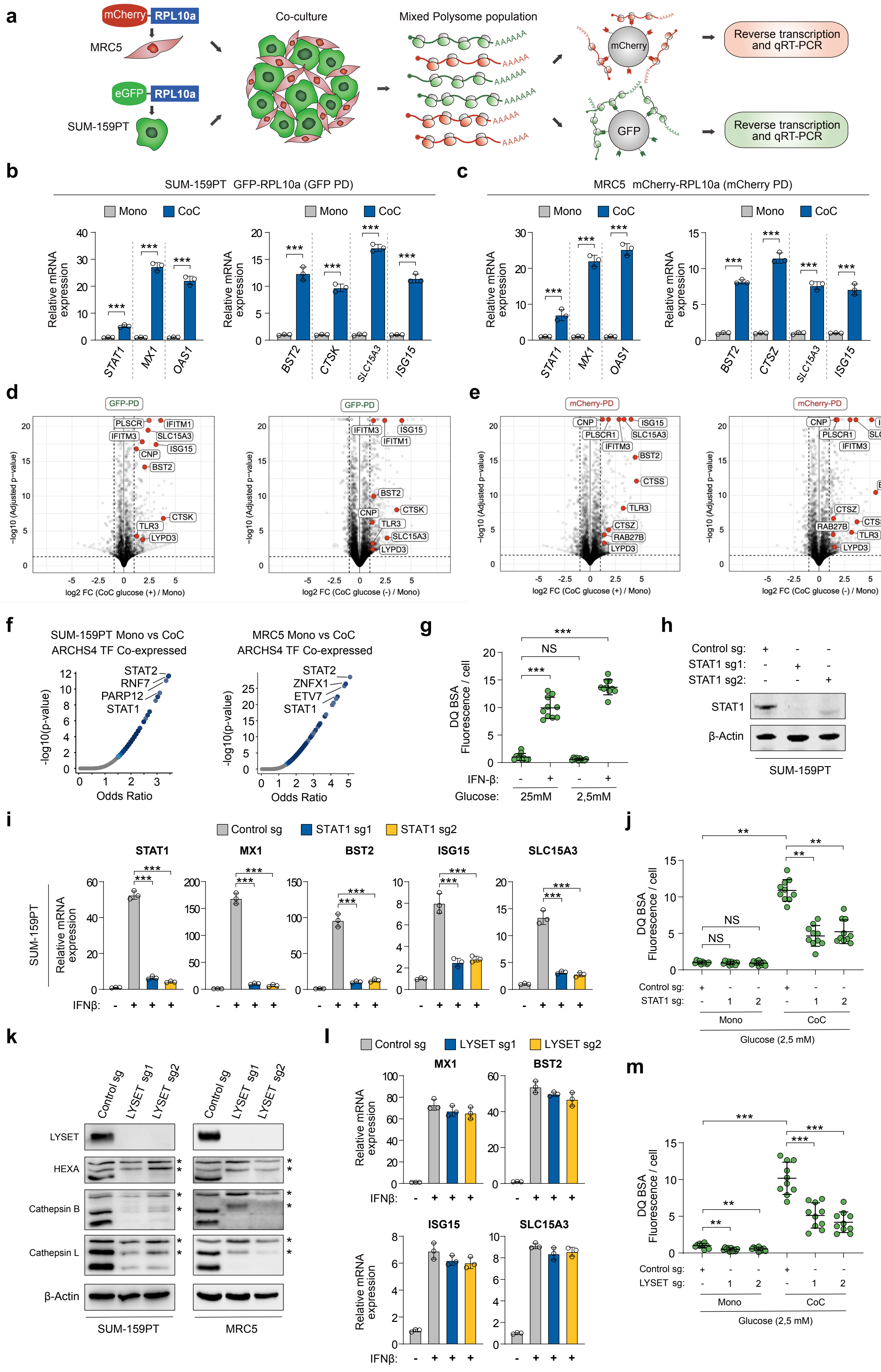

**Supplementary Figure 4. Type I IFN signaling increases lysosomal catabolism upon heterotypic cell interactions.**

**a,** Schematic diagram illustrating the isolation and quantification of polysome-associated transcripts from co-cultures of SUM-159PT-GFP-RPL10a breast cancer cells and MRC5-mCherry-RPL10a fibroblasts.

**b,** qRT-PCR analysis was conducted to measure the expression of the indicated genes in SUM-159PT-GFP-RPL10a cells growing as mono- and co-cultures with MRC5-mCherry-RPL10a fibroblasts. Data represent mean ± SD (n = 3); ***P < 0.001 by Student's t-test. PD, pull-down.

**c,** qRT-PCR assay showing the expression of the indicated genes in MRC5-mCherry-RPL10a fibroblasts growing as mono- and co-cultures with SUM-159PT-GFP-RPL10a cells. Data represent mean ± SD (n = 3); ***P < 0.001 by Student's t-test. PD, pull-down.

**d-e,** Volcano plot illustrating displaying differential expression of genes based on Ribo-seq counts in mono- and co-cultures of SUM-159PT GFP-RPL10a (d) and MRC5 mCherry-RPL10a (e) grown in full ((+), 25mM) or glucose-deprived ((-), 2,5mM) medium. Selected lysosomal genes are highlighted in red.

**f,** Volcano plot illustrating the significance of the upregulated lysosomal gene set in relation to its odds ratio. Every data point represents an individual gene set. The x-axis, represents the odds ratio calculated for each gene set, while the y-axis shows the negative logarithm of the p-value associated with the gene set. Larger data points displayed in blue indicate statistically significant terms (p-value < 0.05), while smaller gray data points represent terms that did not reach statistical significance. The intensity of the blue color of a data point corresponds to its level of significance, with darker blue points indicating higher significance.

**g,** DQ BSA quantification in SUM-159PT cells treated with vehicle or IFN-β (10 ng/mL) for 48hrs. Cells were grown in full medium (Glucose 25mM) or in low-glucose (2,5mM) medium. Data represent mean ± SD (n =10); NS, non-significant; ***P < 0.001 by Student's t-test.

**h,** Western blots on cell extracts from SUM-159PT-GFP-RPL10a cells transduced with sgRNAs targeting STAT1.

**i,** qRT-PCR quantification of the indicated genes in SUM-159PT-GFP-RPL10a transduced with sgRNAs targeting STAT1 and treated with IFN-β (10 ng/mL) for 24hrs. Data represent mean ± SD (n = 3); ***P < 0.001 by Student's t-test.

**j,** Quantification of DQ BSA fluorescence in SUM-159PT GFP-RPL10a transduced with sgRNAs targeting the STAT1 gene or a control sequence growing as mono- or co-cultures (CoC) with MRC5 cells. Cells were grown in low-glucose condition (2,5mM). Data represent mean ± SD (n ≥10); NS, non-significant; **P < 0.01 by Student's t-test.

**k,** Western blot on cell lysates from SUM-159PT-GFP-RPL10a (left panel) and MRC5-mCherry-RPL10a (right panel) cells transduced with sgRNAs targeting LYSET. *Denotes immature lysosomal enzymes.

**l,** qRT-PCR assay of the indicated genes in SUM-159PT GFP-RPL10a transduced with sgRNAs against LYSET and treated with IFNβ (10 ng/mL) for 24hrs. Data represent mean ± SD (n = 3).

**m,** Fluorescence quantification of DQ BSA in SUM-159PT GFP-RPL10a cells transduced with sgRNAs targeting either the LYSET gene or a control sequence, cultured as mono- or co-cultures (CoC) with MRC5 cells. The cells were maintained under low-glucose conditions (2.5 mM). Data are presented as mean ± SD (n =10); **P < 0.01; ***P < 0.001 by Student's t-test.

#### Supplementary Figure 5.

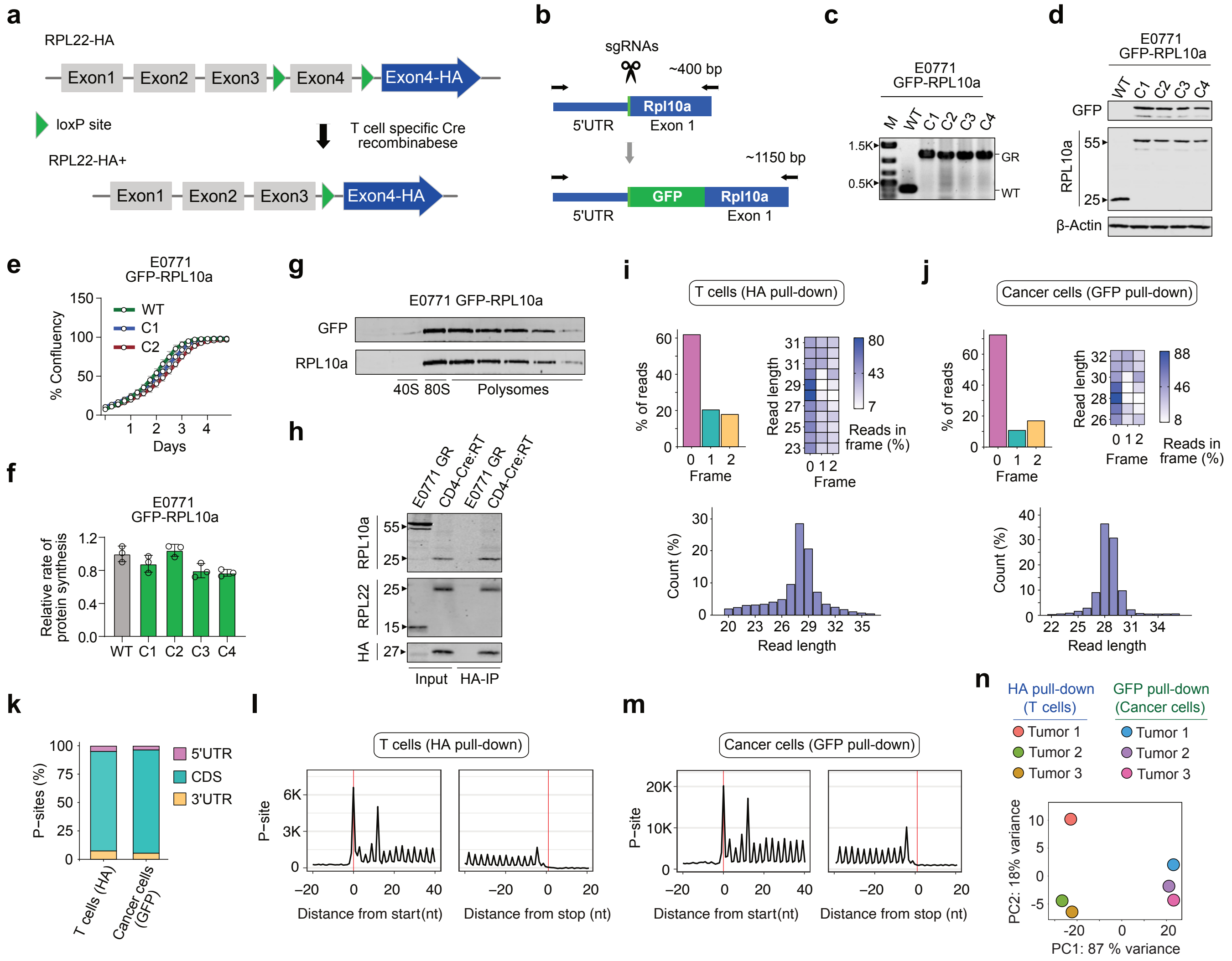

**Supplementary Figure 5. DualRP enables the study of ribosome occupancy in multiple cell compartments of the TME.**

**a,** Schematic diagram of CD4-Cre:RiboTag mice generation. The *Rpl22* gene was subjected to targeted modification through homologous recombination, where loxP sites were introduced to flank the final exon (Exon4). Downstream of the original Exon 4, a new Exon 4 was inserted into the *Rpl22* locus sequence. Activation of the CD4-Cre driver cleaved the loxP sites flanking the initial Exon 4, excising of the floxed exon. Consequently, the Exon4 containing the HA tag becomes incorporated into the RPL22 mRNA, giving rise to the production of RPL22 with an HA tag specifically in T cells (depicted in blue).

**b,** Paired CRISPR–Cas9 nickase approach for tagging the mouse *Rpl10a* gene.

**c,** Gene editing in E0771 cells to incorporate GFP to the N-terminus of *Rpl10a*. Homozygous incorporation was confirmed by PCR.

**d,** Western blot on lysates from clones of E0771 GFP-RPL10a cells, comparing WT or tagged-RPL10a.

**e,** IncuCyte cell proliferation curves of E0771 clones expressing GFP-RPL10a. Data represent mean ± SD (n = 6)

**f,** OP-Puro incorporation assay in E0771 clones expressing GFP-RPL10a. Data represent mean ± SD (n = 3)

**g,** Immunoblot assay was conducted using proteins isolated from individual fractions across the sucrose gradients of cell lysates from E0771 cells expressing GFP-RPL10a.

**h,** Spleens from CD4-Cre:RiboTag (CD4-Cre:RT) mice and E0771 GFP-RPL10a (E0771 GR) cells were lysed and then subjected to immunoprecipitation (IP) using an HA antibody. Both the input and IP fractions were subsequently probed by Western blotting with the specified antibodies.

**i-j,** Frame and read-length distributions of the 5' end of RPFs from ribosome profiling libraries prepared from anti-HA and anti-GFP pull-downs of E0771 tumors.

**k,** Percentage of reads mapping to the 5′ UTR, coding sequence (CDS), and 3′ UTR of mRNAs from data sets generated from anti-HA and anti-GFP pull-downs of E0771 tumors**.**

**l-m,** RPF Coverage around START and STOP codons in ribosome profiling libraries derived from the indicated tumor cell compartments.

**n,** Principal Component Analysis (PCA) from ribosome profiling libraries generated from the indicated pull-downs of E0771 tumors.

#### Supplementary Figure 6.

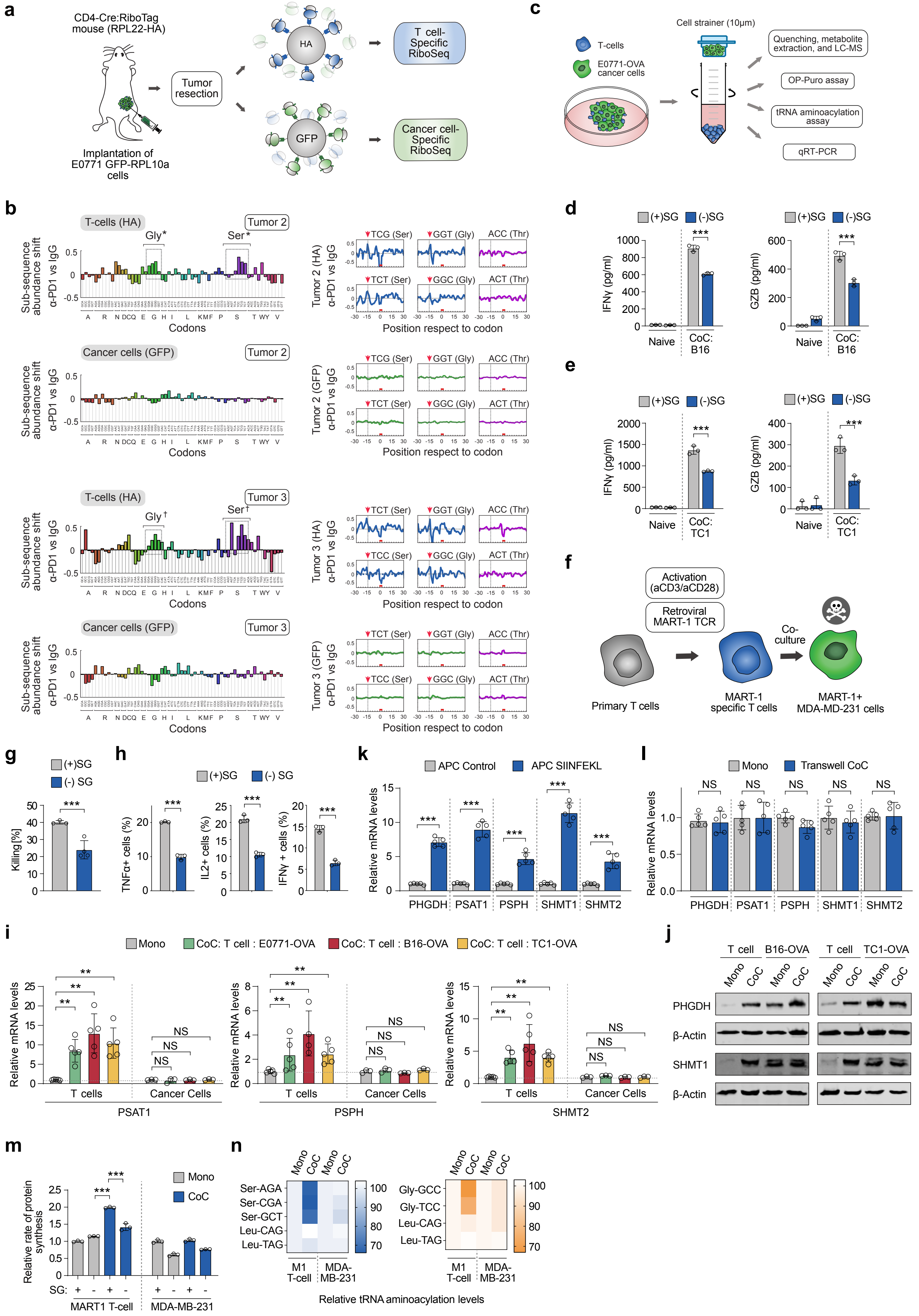

**Supplementary Figure 6. DualRP identifies serine and glycine as crucial nutrients for T cell protein synthesis in the TME during immune checkpoint blockade.**

**a,** Schematic depicting the *in vivo* Dual-RP experiment. Tumors are flash frozen after resection, lysed, and ribosomes from each cell type are immunoprecipitated using either anti-HA- or anti-GFP-coated beads.

**b,** Diricore analysis of independent tumors treated as described in (a). Ribosome stalling in tumor-infiltrated T cells (HA) and E0771 tumor cells (GFP) are shown in the upper and lower panels, respectively. Ribosome density at the A site is shown. * Out-of-frame analysis for tumor 2, $P < 0.01$ for the TCC, TCG, TCT Ser codons and for the GGA, GGC, GGG Gly codons. † Out-of-frame analysis for tumor 3, $P < 0.01$ for the AGT, TCC, TCG, TCT Ser codons and for the GGC, GGG, GGT Gly codons.

**c,** Schematic of the cell-size-based separation of cancer cells and CD8+ T cells.

**d-e,** Quatification of IFNγ and GzmB levels by CD8+ T cells when co-culture with B16-OVA (d) or TC1-OVA (e) cells in full medium or serine/glycine-deprived medium. Data represent mean ± SD (n = 3); ***$P < 0.001$ by Student's t-test.

**f,** Schematic representation of the human co-culture system of antigen-specific T cells and breast cancer cells. T cells were sourced from human peripheral blood mononuclear cells (PBMCs), then activated, and subsequently modified with a MART-1-specific T cell receptor (TCR). Following this, MDA-MD-231 cells expressing HLA-A2 were loaded with MART-1 peptide and co-cultured with the T cells carrying the engineered TCR.

**g-h,** Killing efficiency (g) and quatification of TNFα, IL-2, and IFNγ levels produced by MART-1 T cells (h) when co-cultured with MDA-MD-231 in full medium or serine/glycine-deprived medium. Data represent mean ± SD (n = 3); ***$P < 0.001$ by Student's t-test.

**i-j**, qRT-PCR (i) and Western blot analysis (j) were carried out for the specified genes in co-cultures of OT-I cells and cancer cell lines expressing OVA. Cell separation based on size was performed after 24 hours of co-culture (refer to FigureS 6c). Data represent mean ± SD (n = 5 for T cells and n = 3 for cancer cells). NS, not significant; **$P < 0.01$; by Student's t-test.

**k,** qRT-PCR of the indicated genes in OT-I T cells co-cultured with the antigen presenting cell (APC) line DC 2.4 treated with vehicle (APC control) or pulsed with SIINFEKL peptide (APC SIINFEKL). Data represent mean ± SD (n = 5). ***$P < 0.001$; by Student's t-test.

**l,** qRT-PCR quantification of the indicated genes in OT-I T cells co-cultured with E0771-OVA cells in a transwell plate to prevent direct physical contact. Data represent mean ± SD (n = 5). NS, non-significant by Student's t-test.

**m,** OP-Puro incorporation assay in MART-1 T cells and MDA-MB-231 breast cancer cells growing as mono- or co-cultures. Cell were grown in full medium (SG+) or serine/glycine-deprived medium (SG-). Data represent mean ± SD (n = 3); ***$P < 0.001$ by Student's t-test.

**n,** Aminoacylation assay in MART-1 (M1) T cells and MDA-MB-231 breast cancer cells growing as mono- or co-cultures. Ser-tRNAs, Gly-tRNAs and control Leu-tRNAs were analyzed. Data represent mean ± SD (n=3).

#### Supplementary Figure 7.

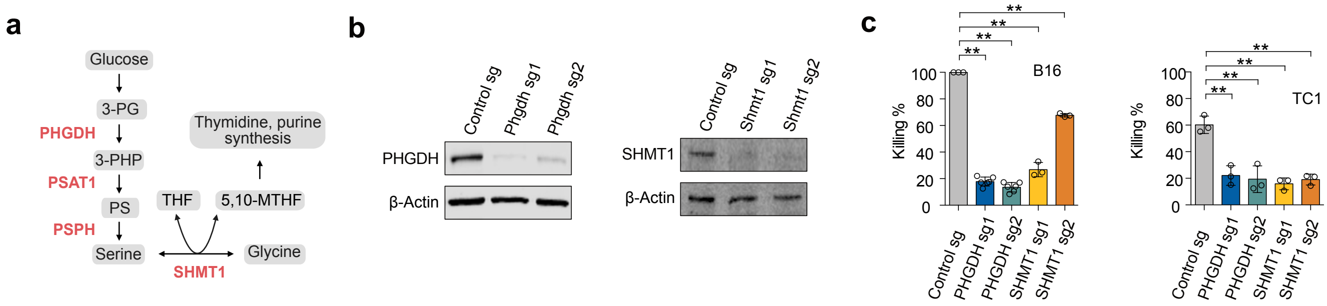

**Supplementary Figure 7. The serine synthesis and one-carbon metabolism pathways are required for efficient cytotoxic activity in CD8$^+$ T cells.**

**a,** Schematic representation of the serine synthesis and the one-carbon metabolism pathway. 3-PG, 3-phosphoglycerate; 3-PHP, 3-phosphohydroxypyruvate; PS, phosphoserine; PHGDH, 3-phosphoglycerate dehydrogenase; PSAT1, phosphoserine amino-transferase1; PSPH, phosphoserine phosphatase; THF, tetrahydrofolate; 5,10-MTHF, 5,10-methylene tetrahydrofolate; SHMT1, serine hydroxymethyltransferase.

**b,** Western blots depicting PHGDH (left panels) or SHMT1 (right panels) protein levels in mouse CD8+ T cells transduced with either control or sgRNAs targeting the Phgdh or Shmt1 genes.

**c,** Cytotoxicity assay was conducted using CD8+ T cells expressing sgRNAs targeting Phgdh or Shmt1. These T cells were co-cultured with pulsed B16 (left panel) or TC1 cells (right panel) for a duration of 24 hours.

#### Supplementary Figure 8.

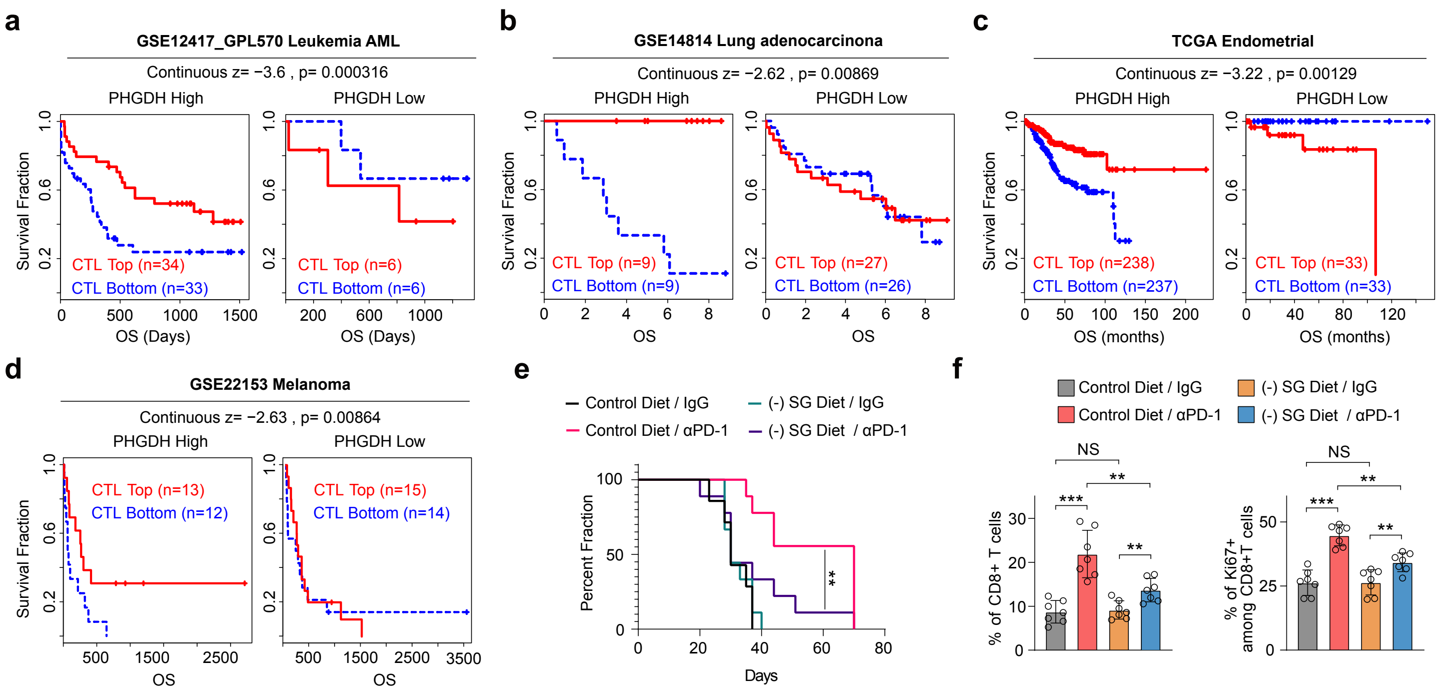

**Supplementary Figure 8. PHGDH predicts cancer immunotherapy outcomes in patients.**
**a-d,** TIDE analyses were conducted on PHGDH expression within T cell dysfunction signatures linked to increased survival in patients with Acute Myeloid Leukemia (AML) (a), Lung Adenocarcinoma (b), Endometrial Cancer (c), and Melanoma (d).
**e,** Survival over time was assessed in mice inoculated with E0771 tumor cells and fed either a control or a (-) SG diet 7 days later. When tumors became palpable, the animals were treated with either IgG (2 μg/μl) or anti-PD1 (2 μg/μl). $^{**}P < 0.01$ by log-rank (Mantel–Cox) test.
**f,** Quantification of the proportion of CD8+ T cells in tumors. Data represent mean ± SD (n = 7). NS, not significant; $^{**}P < 0.01$; $^{**}P < 0.001$; by Student's t-test.
